# Bovine H5N1 influenza viruses have adapted to more efficiently use receptors abundant in cattle

**DOI:** 10.64898/2026.04.02.715584

**Authors:** Jack A. Hassard, Jiayun Yang, Dorothy E. D. P. Hawkins, Jonathan E. Pekar, Bernadeta Dadonaite, Jin Yu, Samuel A. S. Richardson, Rute M. Pinto, Kristel Ramirez Valdez, Philippe Lemey, Jessica L. Quantrill, Chris Neil, Lorin Adams, Jinghan Xue, Natacha S. Ogando, Tereza Masonou, Katie-Marie Case, Jila Ajeian, Maximillian N. J. Woodall, Rebecca A. Ross, Nicholas Hudson, Kan Zhong, Hongzhi Cao, Samuel Jones, Hannah Klim, Brian R. Wasik, Desi N. Dermawan, Jean-Remy Sadeyen, Dirk Werling, Dylan Yaffy, Joe James, Alejandro Nunez, Paul Digard, Ian H. Brown, Daniel H. Goldhill, Pablo R. Murcia, Claire M. Smith, Yan Liu, Jesse D. Bloom, Munir Iqbal, Wendy S. Barclay, Stuart M. Haslam, Thomas P. Peacock

**Author notes:** co-first authors (Jack A. Hassard, Jiayun Yang, Dorothy E. D. P. Hawkins, Jonathan E. Pekar).

## Abstract

Sustained mammal-to-mammal transmission of high pathogenicity H5N1 avian influenza viruses is reshaping the host range of these pathogens. One of the longest-running mammalian transmission chains involves the B3.13 genotype circulating in U.S. dairy cattle which was detected in early 2024. Genomic analyses revealed selection and rapid fixation of haemagglutinin mutations D104G and V147M. We demonstrate, via glycomic profiling, that bovine tissues, including the mammary gland, are enriched in N- and O-linked glycans capped with N-glycolylneuraminic acid (NeuGc), a sialic acid which is not synthesised by humans and birds, which instead express only N-acetylneuraminic acid (NeuAc). Early cattle H5 viruses poorly recognised NeuGc, but D104G and V147M enabled efficient engagement of both NeuAc- and NeuGc-containing receptors. These mutations enhanced replication in bovine mammary tissue without major attenuation of replication in human lung and primary nasal epithelial cells. NeuGc-driven receptor adaptation therefore promotes viral fitness in cattle without majorly changing zoonotic risk. Deep mutational scanning was able to identify alternative haemagglutinin substitutions that confer NeuGc usage and represent surveillance markers for emerging cattle H5 lineages. Finally, structural analyses suggest a potential molecular mechanism for the dual binding of the cattle-adapted viruses.

## Introduction

In 2020, a novel genotype of clade 2.3.4.4b H5N1 influenza virus arose and began spreading around the world, triggering a panzootic^1^. This virus was initially associated with wild waterfowl and seabirds but was later marked by a high frequency of spillover into mammalian hosts^2^. While most spillovers into mammals were dead-end or transient, several examples were found with strong support for sustained mammal-to-mammal transmission^2^. One of the longer lasting mammal clusters is in US dairy cattle. In early 2024, a mystery milk drop syndrome was identified as being caused by H5N1 infection^3,4^. Relative to the highly severe H5N1 infections seen in carnivores, this infection was mild and thought to be spread mainly within farms by contaminated milking machinery or between farms/states by movement of infected animals or equipment^4^.

Most influenza A viruses use glycans terminating in sialic acid as their primary receptor. The specific and dynamic display of these sialylated glycans differ between species. For example, the human respiratory tract is abundant in glycans terminating in α2-6-linked sialic acid^5^, whilst the avian gastrointestinal tract, the major site of replication for avian influenza viruses, contains a high abundance of α2-3-linked sialic acid. Consequently, human seasonal and pandemic influenza viruses preferentially bind to α2-6-linked sialic acid and avian influenza viruses, including H5N1 viruses, generally have a strong preference towards α2-3-linked sialic acids^6,7^.

Beyond differences in the linkage to the penultimate sugar, the type of sialic acid can also differ between species. Humans and birds predominantly contain N-acetylneuraminic acid (NeuAc), which is usable by influenza viruses^5,8–10^, however, many mammalian species contain a second type of sialic acid, N-glycolylneuraminic acid (NeuGc). During glycan biosynthesis, NeuAc can be modified by hydroxylation of the N-acetyl on carbon 5 to N-glycolyl by the enzyme Cytidine Monophospho-N-Acetylneuraminic Acid Hydroxylase (CMAH) to form NeuGc. Alongside humans and birds, several important influenza hosts, including mustelids^11^, seals^12^ and Western dog breeds^13^, only synthesise the NeuAc form of sialic acid. This is due to these species lacking functional CMAH, either due to low expression (such as in Western dog breeds), or a cmah gene that is pseudogenised (such as in humans) or entirely deleted (such as in birds, mustelids and seals). In other influenza host species, such as pigs^14,15^ and horses^16^, high expression of CMAH in tissues results in the display of NeuGc on glycans. Most influenza viruses preferentially bind to sialylated glycans terminating in NeuAc, and, at best, poorly use NeuGc-containing glycans as a receptor^17^. However, a now extinct equine H7N7 virus showed a strong preference towards NeuGc^17,18^.

To date, multiple studies have used both immunohistochemical and immunofluorescence techniques to probe influenza virus receptor distribution in cattle tissues using lectins and influenza haemagglutinins (HAs)^7,19–21^. Generally, these analyses have shown that certain H5 HAs, such as those from the clade 2.3.4.4b viruses, are able to bind to cells in the mammary gland, but not throughout the respiratory tract. It has been suggested that modified sialic acids which are not conventional influenza virus receptors, such as NeuGc, are more abundant in the bovine respiratory tract^19^. However, to date, no detailed glycomic characterisation of cattle tissues has been undertaken.

Adaptation of avian influenza viruses to mammalian hosts is required for sustained transmission. We and others have previously described how early adaptations in the polymerase genes of bovine H5N1 allow the virus to better infect and replicate in diverse mammalian species^22–24^. During its evolution in cattle, several HA mutations have arisen, and in some cases fixed, in bovine H5N1. Given these do not appear to impact α2-6 NeuAc binding of these viruses^6^, we therefore undertook a detailed investigation into the impact of these mutations on binding to structurally defined glycans present in cattle, and what impact these mutations have on replication in human cells. By linking host glycan landscapes to viral receptor adaptation, we reveal a general mechanism by which influenza viruses can evolve following their emergence in mammalian hosts.

## Results

### Several Haemagglutinin mutations are positively selected for and have reached fixation in cattle H5N1 sequences

To investigate whether there was selection within the HA gene of B3.13 genotypes infecting cattle over its approximately first 2 years of evolution (Figure 1A), we examined ∼4000 virus genomes sampled from cattle and humans. We found multiple residues with evidence of positive selection (Supplementary Figure S1, Supplementary Data S1, Supplementary Table 1), including D104G and V147M (Figure 1B, C; H5 immature numbering used throughout, equivalent to D88G and V131M in mature H5 numbering, see supplementary Table S1 for conversion). V147M arose at least 8 times independently, before fixation in recent cattle H5N1 viruses alongside D104G (Figure 1D). All 2026 sequences (as of September 2026) continue to contain both mutations. Structurally V147M lies within the 130-loop, while D104G lies below the 130-loop (Supplementary Figure S1). From previous deep mutational scanning data in a North American clade 2.3.4.4b HA neither mutation is predicted to influence binding to α2-6 sialylated glycans^25^. Although residues within the 130-loop are not often described as influencing α2-3 versus α2-6 binding, they directly interact with the leading edge of the sialic acid moiety and have previously been described as contributing to binding modified sialic acids^26^. To explore this possibility, we performed an in-depth analysis of the types of sialylated glycans found in different cattle tissues.

**Figure 1.**
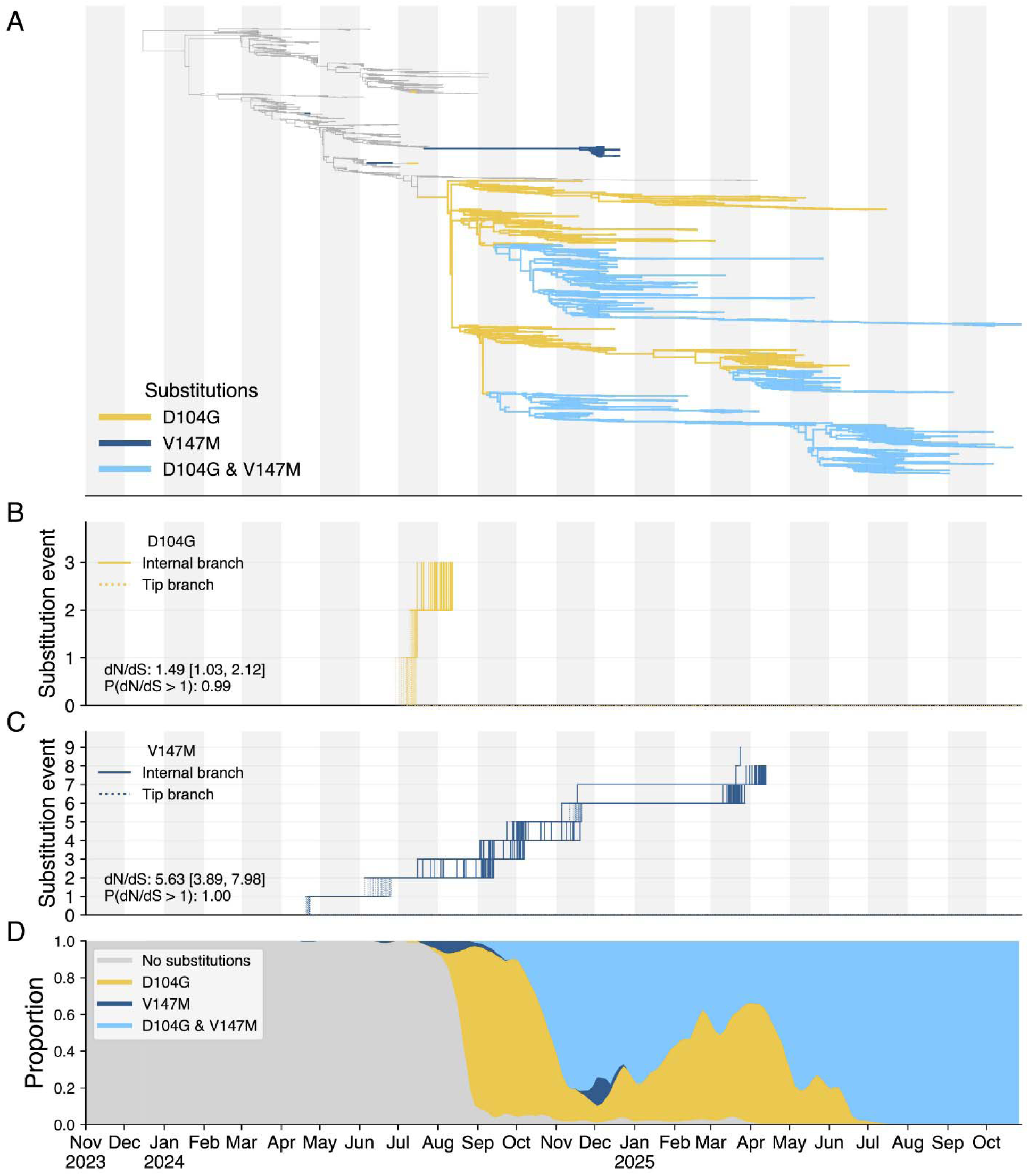
Positively selected mutations in the cattle H5N1 head domain have reached fixation as the virus continues to circulate. **A.** Time-scaled phylogenetic tree of 3,929 H5N1 sequences sampled from cattle and humans, with amino acid substitutions D104G and V147M indicated in yellow and dark blue, respectively, when only one is present, and in light blue when both are present (see Methods). The repeated emergence of D104G (**B**) and V147M (**C**) across internal branches (solid lines) and tip branches (dotted lines). Each stepwise path corresponds to the evolutionary trajectory of these substitutions in one posterior sample (n=322). The dN/dS value (median and 95% highest posterior density [HPD]) and its probability of being above 1 are indicated in the bottom left of the panels. **D.** The proportion of lineages with either D104G, V147M, neither substitution, or both substitutions based on the highest independent posterior subtree reconstruction of H5N1.

### The bovine mammary gland and respiratory tract contain an abundance of NeuGc sialylated glycans

We characterised the types of sialylated N- and O-linked glycans in cattle from four different relevant tissues: mammary gland, teat, trachea and lung. Glycomic profiling of the bovine tissues (Figure 2B-E, Supplementary figures S2-S5, Supplementary Tables 3 and 4) revealed the presence of a diverse range of complex N-glycans terminating in sialic acid. Both NeuAc- (e.g. m/z 2605, 2792, 2966), and NeuGc- (e.g. m/z 2635, 2852, 3026) or a mixture of NeuAc- and NeuGc- (e.g. m/z 2822, 2996) capped N-glycans were observed across all tissues tested. Glycans ranging from bi-antennary with no sialic acid residues (e.g. m/z 2244) to tetra-antennary with up to 5 sialic acid residues were detected (e.g. m/z 5213, 5243, 5662). Glycans expressing 2 sialic acid residues on a single branch (e.g. m/z 3154 3184, 3214) were confirmed to express the additional sialic acid on the GlcNAc by tandem MS/MS analysis (Supplementary figure S4). Glycans with up to 7 LacNAc units were detected (e.g. m/z 5621) suggesting the presence of extended poly-LacNAc chains and I-branches which could be capped with sialic acid. Relative abundances of specific glycans varied between the tissues, with differences in the abundance of glycans capped with NeuAc, NeuGc, or Galα1-3Gal.

**Figure 2:**
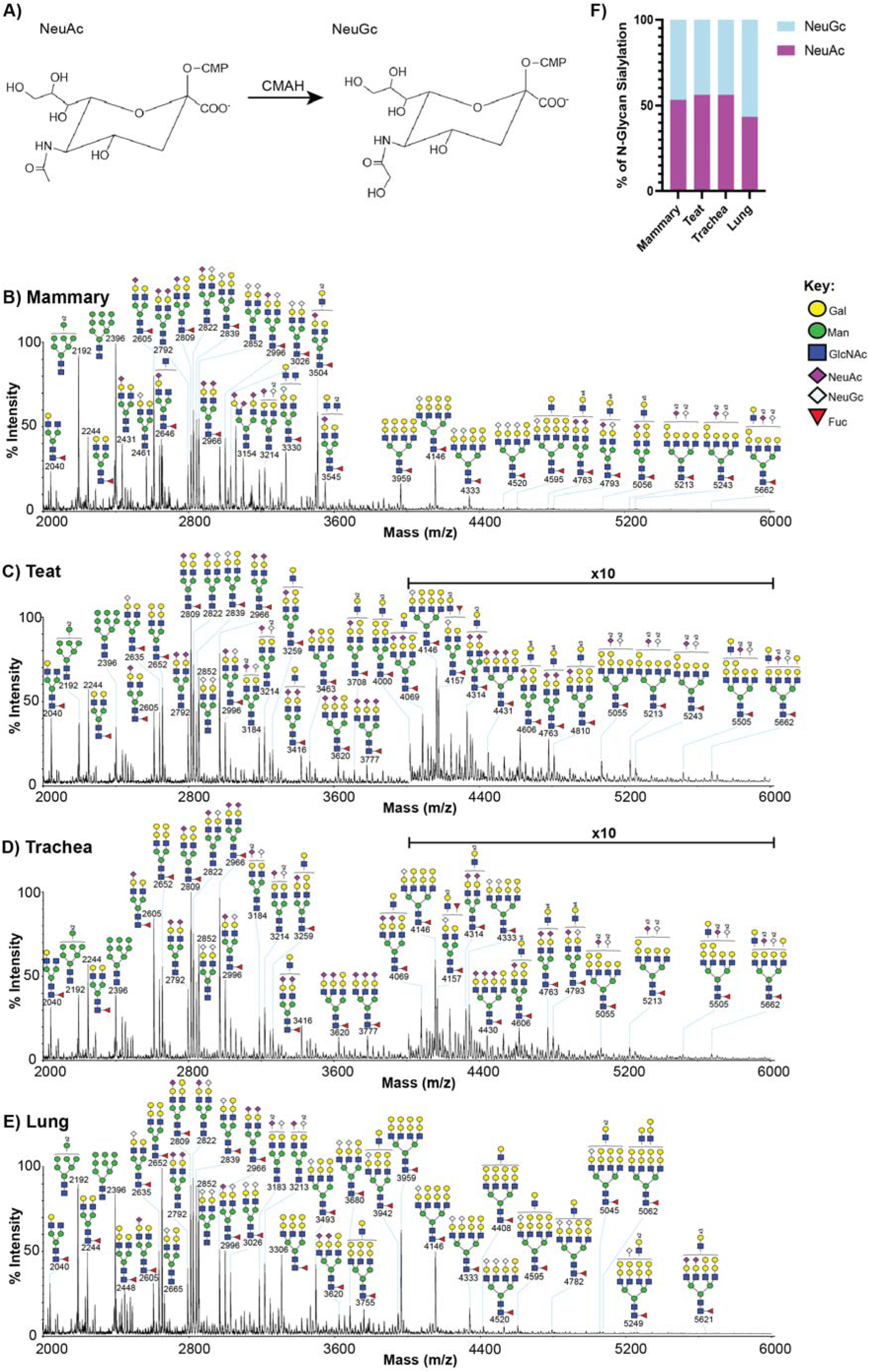
Bovine tissues contain glycans terminating in a mixture of NeuAc and NeuGc. **A.** schematic of conversion of CMP-NeuAc into CMP-NeuGc by CMAH. Annotated MALDI-TOF spectra of bovine mammary **(B)**, teat **(C)**, trachea **(D)** and lung **(E)** tissue N-glycans from a lactating cattle. Annotations show [M+Na]^+^ molecular ions. Peak annotation is based on composition, biosynthetic knowledge and MS/MS analysis. **F.** Calculated proportions of NeuAc and NeuGc in bovine samples. Data shown is representative of N = 4 for mammary and teat samples, and N = 2 for tracheal and lung samples.

Glycan distribution was comparable between cows, following normalization of the glycan spectra we calculated that the ratios of NeuAc to NeuGc in the tissues tested were approximately equal (Fig. 2F), in tissues from both lactating and non-lactating animals (Supplementary Figure S3). The proportions of α2-3 and α2-6 linkage configurations were also calculated following treatment of the glycans with Sialidase-S, which specifically cleaves α2-3 linked sialic acid (Supplemental figures S3-17). We determined that α2-3 linked sialic acid was moderately more abundant for the tissues tested, both for NeuAc and NeuGc sialic acid types.

Analysis of bovine O-glycans from the tissues showed core 1, core 2, core 3 and core 4 structures which were often extended and capped with sialic acid (Supplementary figure S2). Both NeuAc and NeuGc capped structures were present however the majority of sialylation was NeuAc and α2-3 linked in the O-glycans.

Overall, we demonstrate that bovine mammary and respiratory tissues contain a high abundance of α2-3 linked NeuAc, the preferred receptor type for early cattle H5N1 isolates and consistent with other studies^6,7,27,28^. However, due to the near equal abundance of α2-3 linked NeuGc containing N-glycans we hypothesised this receptor type could be driving mutations in the bovine H5N1 HA that had been selected for over the months the virus had been circulating in cattle^6^.

### Prevalent mutations in the haemagglutinin of bovine H5N1 allow the virus to better bind and utilise NeuGc-containing glycans abundant in the cattle

Upon establishing that cattle tissues were abundant in sialylated glycans with terminal NeuGc, we generated a panel of H5N1 mutants in the background of a reverse genetics-derived cattle/Texas HA, with the polybasic cleavage site removed and combined with the internal gene segments (segments 1-3, 5 and 7-8) of the attenuated, laboratory-adapted PR8 strain, as previously described^6^. We chose mutations previously found to be under positive selection, that had arisen in cattle viruses, and that were proximal to the receptor binding site. Furthermore, we generated a Y173A mutant as a positive control, which has previously been described as switching H5 HA from a preference for NeuAc to NeuGc^17,29^. These mutants were rescued, propagated in embryonated hens’ eggs, sequenced, then purified by ultracentrifugation on a sucrose gradient.

Using these purified viruses we tested receptor binding using bio-layer interferometry (BLI), which directly measures the kinetics of virus binding to receptor analogues, and glycan microarrays against a panel of α2-3 NeuAc and NeuGc containing glycans (Supplementary figures S18-S21). We found the early cattle H5N1 HA (cattle/Texas WT) showed a strong preference for an α2-3-linked NeuAc-containing receptor analogue over the equivalent analogues containing NeuGc, and no detectable binding to analogues with either NeuAc or NeuGc attached via an α2-6-linkage (Figure 3A-C). Conversely, cattle/California, a more recent cattle H5N1 virus containing the combination of HA mutations D104G, S110N, and V147M, strongly bound 3SLN(NeuGc) and was shown to bind NeuGc containing analogues to a similar level as the NeuAc equivalent (Figure 3D). Testing individual mutants in the cattle/Texas genetic background we mapped this switch to V147M and to a lesser extent D104G (Figure 3E-G). The control mutant, Y173A, resulted in a switch in preference from 3SLN(NeuAc) to 3SLN(NeuGc) (Figure 3H), as previously described for other H5 viruses^28^. Several further mutations also somewhat enhanced 3SLN(NeuGc) binding including the common early mutation T143A, and some less common mutations such as V147E, V147A, Q154L, and D171N (Supplementary Figures S22-24). No H5N1 mutants tested had detectable binding to either 6SLN(NeuAc) or 6SLN(NeuGc) when probed by BLI.

**Figure 3.**
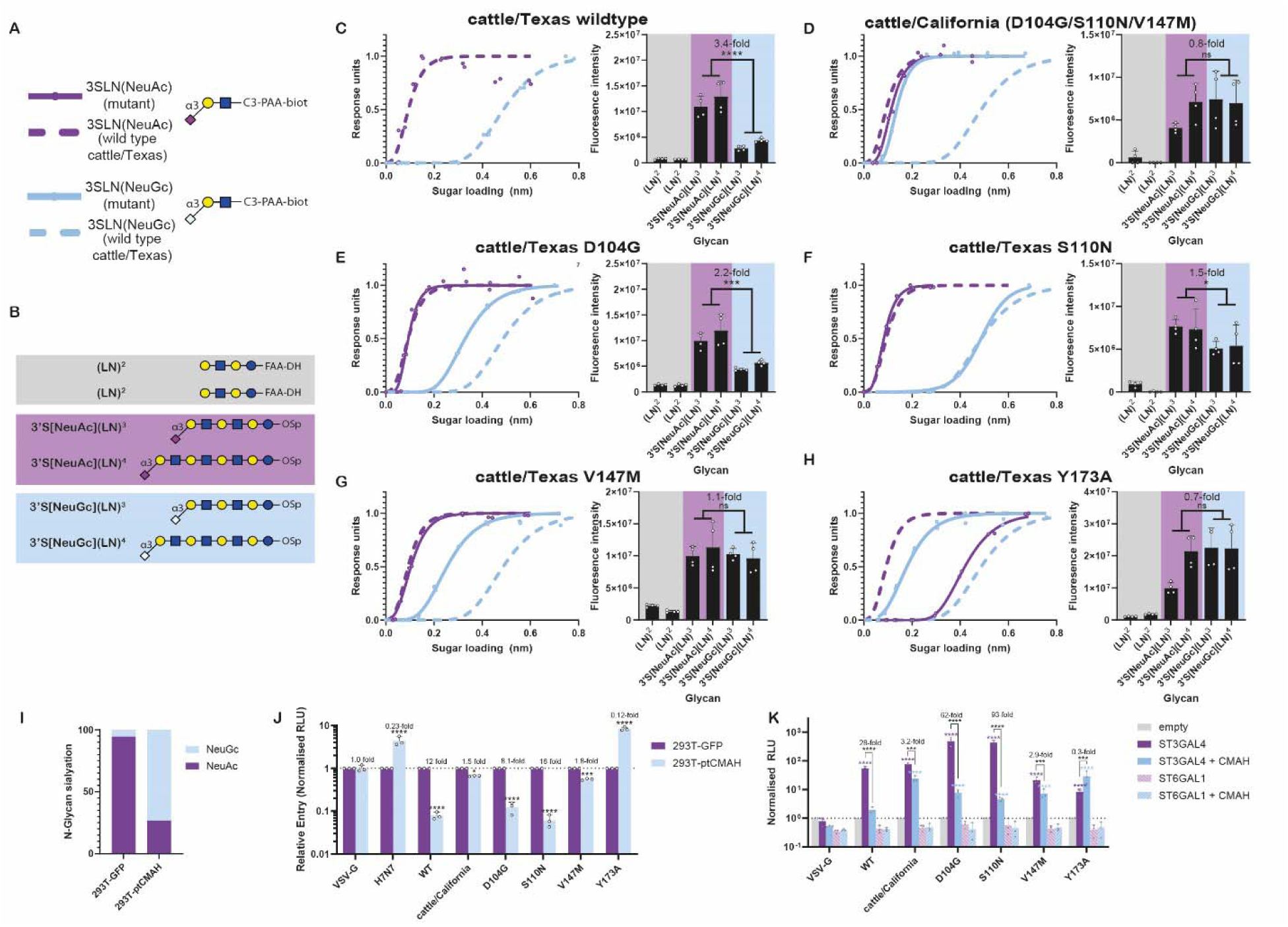
Mutations common in cattle H5N1 viruses enhance the ability to bind receptors containing NeuGc, and increase entry into cells expressing predominantly NeuGc. **A, C-H, left panels.** Bio-layer interferometry plots showing binding of purified viruses to different receptor analogues shown in panel **A.** Solid (named virus/mutant) and dashed (wildtype cattle/Texas), pink [(3SLN(NeuAc)] and pale blue [3SLN(NeuGc)] lines indicate receptor binding to analogues. Data points shown as duplicate repeats with a asymmetric Sigmoidal 5PL curve fitted, constrained to 0 and 1 at the top and bottom. **B, C-H, right panels**. Binding of purified viruses to a panel of receptor analogues (shown in panel **B**) immobilized to glycan arrays. Non-sialylated analogues backed in grey, analogues containing NeuAc backed in pink and NeuGc backed in pale blue. Data shown as a representative N = 2 biological repeats including N = 2 technical repeats (individual biological repeats on an extended library shown in figures S17-18). Statistics performed by two-way ANOVAs with individual variance for each group – e.g. comparing each matched set of analogues containing either NeuAc or NeuGc. **I.** Summary of glycomics data showing the ratio of NeuAc to NeuGc in 293T cells N-glycans overexpressing either GFP as a control, or chimpanzee CMAH. **J.** Relative entry of pseudoviruses carrying the named glycoproteins into 293Ts stably expressing GFP or chimpanzee CMAH. Data normalised to entry into 293T-GFP cells. Data plotted as mean + S.D. with the mean values from N = 3 independent repeats plotted. Statistics performed using 2-way ANOVA with multiple comparisons upon log-transformed data. **K.** Relative entry of pseudoviruses into 293 cells that have been genetically ablated for sialyl-transferases, then transiently transfected with either empty vector and an expression plasmid for sialyl-transferases (ST3GAL4 or ST6GAL1) with or without chimpanzee CMAH. Data plotted as mean + S.D. with the mean values from N = 3 independent repeats plotted. Statistics performed using 2-way ANOVA with multiple comparisons upon log-transformed data. Only differences above empty vector, and between ST3GAL4 and ST3GAL4+CMAH shown. Log-normality throughout determined by Shapiro-Wilk test and QQ plot. Significance shown by asterisks indicating: *, 0.05 ≥ P > 0.01; **, 0.01 ≥ P > 0.001; ***, 0.001 ≥ P > 0.0001; ****, P ≤ 0.0001.

To confirm these biophysical binding results, we used a functional assay employing matched pseudovirus mutants and tested HA-mediated entry into 293T cells either overexpressing GFP (control cells) or chimpanzee CMAH^30^. Glycomic analysis confirmed that stably expressing chimpanzee CMAH skewed these cells towards expressing a high proportion of NeuGc-containing sialylated glycans (Figure 3I, Supplementary Table 5). A pseudovirus containing the HA from cattle/Texas had attenuated entry into 293T-CMAH cells (∼12-fold lower), but this was almost completely overcome by the introduction of D104G or V147M alone or in combination (∼1.5 – lower entry). As controls we included pseudoviruses expressing the HAs of the NeuGc-preferring equine H7 and cattle/Texas Y173A and found these both entered 293T-CMAH cells more efficiently than the control cells. Additionally, we used 293 cells with sialyl-transferases knocked out^31^, and transiently re-expressed either an α2-3- or α2-6-specific sialyltranferase (ST3GAL4 or ST6GAL1, respectively), in combination with excess of either chimpanzee CMAH or empty vector (Figure 3K, Supplementary Figure S25). These data were consistent with those obtained using the stable cell lines and supported the BLI results in that additional binding to NeuGc was only present when an α2-3 sialyltranferase was expressed, with no detectable boost in entry when the α2-6 sialyltranferase was expressed, with or without CMAH overexpression (Figure 3K).

Overall, we have shown using three different methodologies that mutations selected for and fixed in H5N1 during its circulation in cattle have enhanced the virus’s ability to bind and enter via NeuGc-containing glycans abundant in cattle tissues, therefore we next tested if this also had an impact on virus replication in relevant model systems.

### Common cattle H5N1 mutations enhance replication in bovine mammary explants, but attenuate replication in human airway cells

To determine if enhanced NeuGc binding correlated with enhanced replication in a relevant bovine system, we tested the replication of the previously generated 2:6 (H5N1:PR8) viruses in bovine ex vivo mammary explants (Figure 4A). We found that mutants with enhanced NeuGc binding were associated with higher infectious titres, with several reaching significance across multiple time points. Conversely, we found in human Calu-3 lung cells that these mutants showed similar titres through the time course (Figure 4B). We next generated full H5N1 viruses with intact polybasic cleavage sites, and all 8 gene segments from cattle/Texas and tested their replication in primary human nasal epithelial cells (hNECs) derived from three independent donors, differentiated and maintained at air-liquid interface. Consistent with the Calu-3 data, we found that all the tested mutants replicated similarly, or moderately worse than wild type cattle/Texas, significantly so at some time points in some donors (Figure 4C-E).

**Figure 4.**
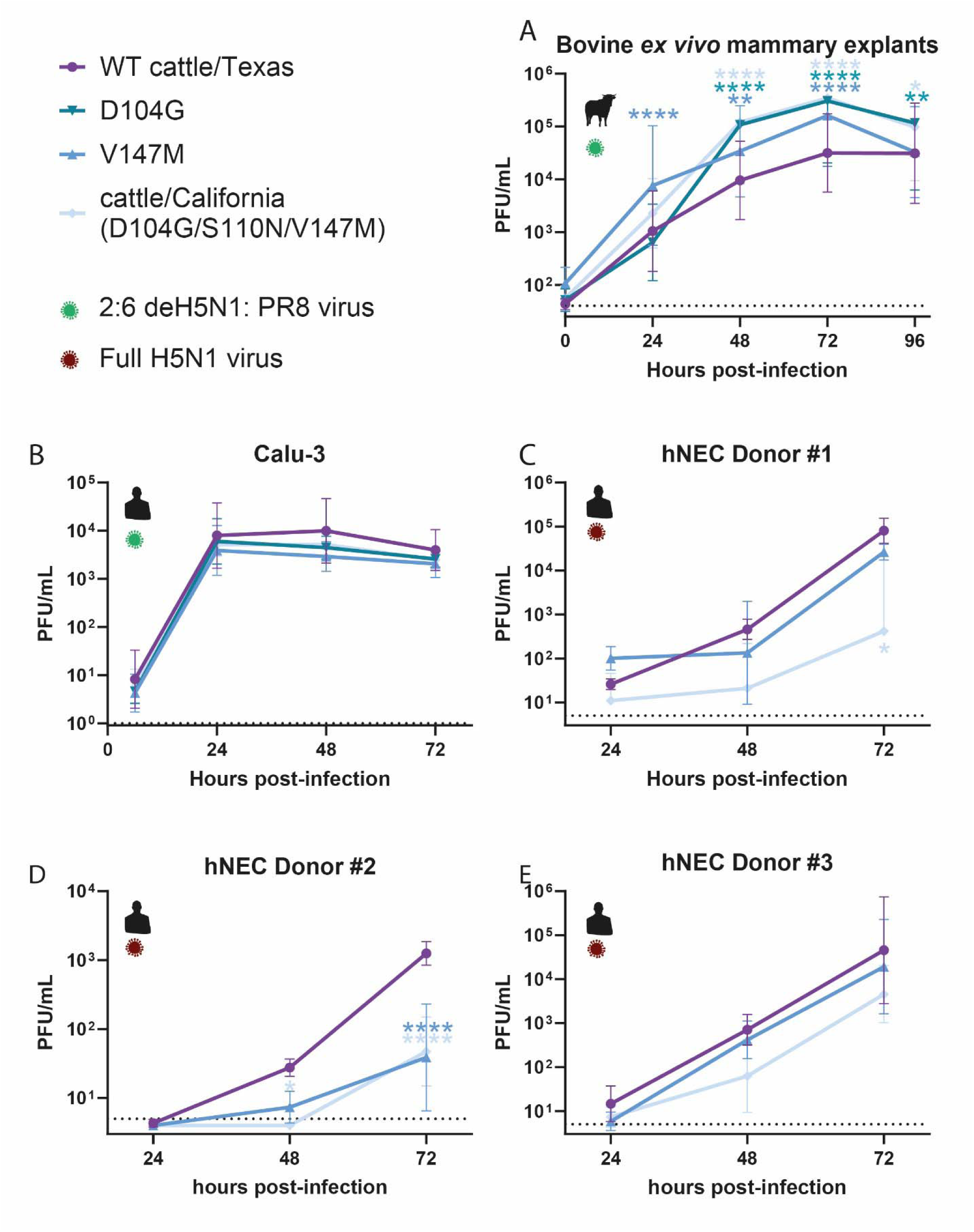
NeuGc-adaptive mutations enhanced replication in cattle explants but neutral or modestly attenuating in human systems. Replication of 2:6 (de-engineered HA and NA from cattle/Texas, remaining genes from PR8) in **A.** bovine mammary explants, from N = 4 donors and N = 52 total repeats (N = 4 from donor 1, N = 12 from donor 2, N = 20 from donor 3, N = 16 from donor 4) and **B.** human calu-3 cells (N = 3 repeats). **C, D, E.** Replication of full isogenic H5N1 viruses in the cattle/Texas background in human primary nasal cultures (hNECs), maintained at air-liquid interface. Data plotted from 3 independent donors and as N=3 technical repeats. Data throughout plotted as mean ± SD. Statistics throughout performed by two-way ANOVA on log-transformed data with multiple comparisons between the wild type and mutants. Significance denoted by asterisks indicating: *, 0.05 ≥ P > 0.01; **, 0.01 ≥ P > 0.001; ***, 0.001 ≥ P > 0.0001; ****, P ≤ 0.0001.

Overall, these data suggest that while these NeuGc binding mutations allow enhanced replication in bovine tissues they are either neutral, or modestly attenuated in their ability to cause zoonotic infection in humans.

### Deep mutational scanning identifies regions of the HA gene associated with enhanced NeuGc usage

Most contemporary H5N1 cattle sequences of the B3.13 genotype contain the combination of D104G and V147M (Figure 1D). However, additional spillovers from wildfowl into cattle, such as the three independent 2025 spillovers of genotype D1.1 into cattle identified in Arizona, Nevada and Wisconsin^32^, currently lack any equivalent overt NeuGc adaptation. We therefore wanted to predict whether alternative routes to gaining NeuGc binding could be occurring or could arise in the future. To do this we utilised a previously generated H5 HA pseudovirus-based deep mutational scanning (DMS) library^25^, which allows the measurement of the impact of almost all possible mutations in HA on entry into cells expressing mainly NeuGc or NeuAc containing-sialic acids.

First, we compared the effects of mutations on entry in control HEK293 cells versus HEK293 cells overexpressing chimpanzee CMAH^30^ (Figure 5A). We identified many mutations that are beneficial for entry in CMAH overexpressing cells but deleterious for entry in HEK293 cells, suggesting these mutations increase NeuGc binding at the expense of binding to NeuAc. (Figure 5B-C, Supplementary Figure S26). When mapped onto a structure of H5N1 HA these mutations largely clustered around the distal end of the sialic acid moiety, close to or directly interacting with where the additional protruding hydroxyl group of NeuGc is expected to lie (Figure 5B), in a similar manner to that described structurally for Y173A^17^. Sites in the 130- and 150-loops, at positions 137-147, 159-160, 166, 168, 173 and 176 in particular showed large increases in entry into the CMAH cells relative to the control cells (Figure 5B-C).

**Figure 5.**
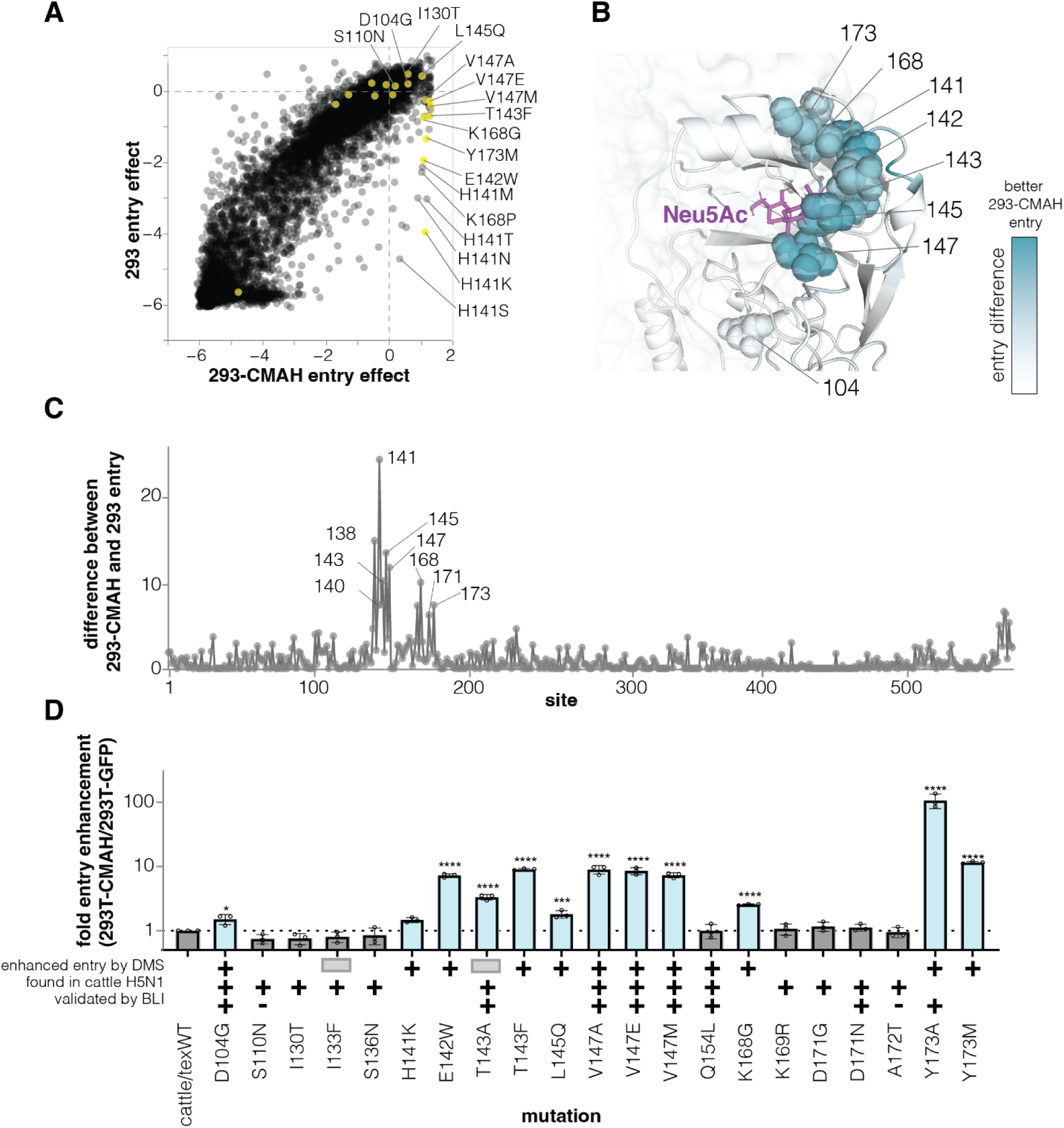
Deep mutational scanning identifies hotspots in the H5 HA associated with enhanced binding to NeuGc containing glycans. **A.** Cell entry effects of mutations on HEK293 (expressing NeuAc) versus HEK293-CMAH cells (expressing NeuGc). Entry effects above 0 indicate mutations that are positive for entry and those below 0 indicate mutations deleterious for entry. Mutations that were chosen for validation in **D** are highlighted in yellow. **B.** Structure of receptor binding pocket in HA with cell entry difference (cell entry effect measured on HEK293 cells subtracted from entry effect on HEK293-CMAH cells) measured by deep mutational scanning (DMS) shown as a heatmap on the structure and bound NeuAc is shown in purple. **C.** Total cell entry difference between HEK293 and HEK293-CMAH cells for every position in HA. Interactive version of this plot as well as mutation-level effects can be viewed at https://dms-vep.org/Flu_H5_American-Wigeon_South-Carolina_2021-H5N1_DMS/NeuGc_usage.html **a** H5 HA pseudovirus, generated in the background of cattle/Texas H5 HA, entry enhancement on HEK293T-CMAH cells compared to HEK293T. Bars in light blue indicate mutations that increase entry on NeuGc expressing cells. Mutations that are measured to increase NeuGc by DMS are indicated by a plus sign; mutations that were not measured in the DMS experiments are indicated by grey rectangles. Mutations found in circulating cattle clade H5 sequences are indicated by plus sign. Mutations that were found to increase or decrease binding to NeuGc by BLI are indicated with plus and minus signs, respectively. Data plotted as the mean of N = 3 independent repeats and normalised to wild-type cattle/Texas. Data plotted as mean +/− S.D. Statistics performed on log-transformed data using one-way ANOVA with multiple comparisons against wild type cattle/Texas. Log-normality determined by Shapiro-Wilk test and QQ plot. Significance shown by asterisks indicating: *, 0.05 ≥ P > 0.01; **, 0.01 ≥ P > 0.001; ***, 0.001 ≥ P > 0.0001; ****, P ≤ 0.0001.

To validate the impact of mutations identified by DMS to increase NeuGc usage, as well as the naturally occurring mutations which were tested by BLI, we generated H5 HA mutants in a lentivirus-based pseudovirus system and tested entry into HEK293T versus HEK293T-CMAH cells (Figure 5D). Sites predicted to increase NeuGc usage showed marked enhancement of entry into HEK293T-CMAH cells relative to HEK293T cells, including mutations D104G, H141K, E142W, T143A/F, L145Q, V147A/E/M, K168G and Y173M. These sites bear particularly close monitoring in cases of spillover of H5N1 into NeuGc containing hosts such as cattle and horses.

### Haemagglutinin receptor binding site flexibility may promoter NeuGc binding

Finally, to understand the mechanistic basis for the dual receptor binding by the cattle/California virus, we determined the structures of cattle/California and cattle/Texas HA ectodomains in the apo and 3SLN(NeuGc) -bound state by single particle cryo-electron microscopy (Figure 6, Supplementary Figure S27). We were unable to resolve any gross structural changes that would explain the difference in binding to NeuGc between cattle/California and cattle/Texas. However, NeuGc binding has previously been associated with increased flexibility of the 130 loop^17^. Position 147 lies within this loop, proximal to the additional hydroxyl group in NeuGc relative to NeuAc, while 104 sits below the 130 loop. We note that valine (cattle/Texas 147) has particularly constrained rotational flexibility due to beta-branching. The previous bio-layer interferometry and DMS data suggest that most amino acid substitutions from V147, including alanine and glutamate which exhibit marked different biochemical properties, also enhance NeuGc binding. Taken together this suggests these other mutations away from the beta-branched valine may also enhance 130-loop flexibility, allowing for greater sialic acid binding promiscuity.

**Figure 6.**
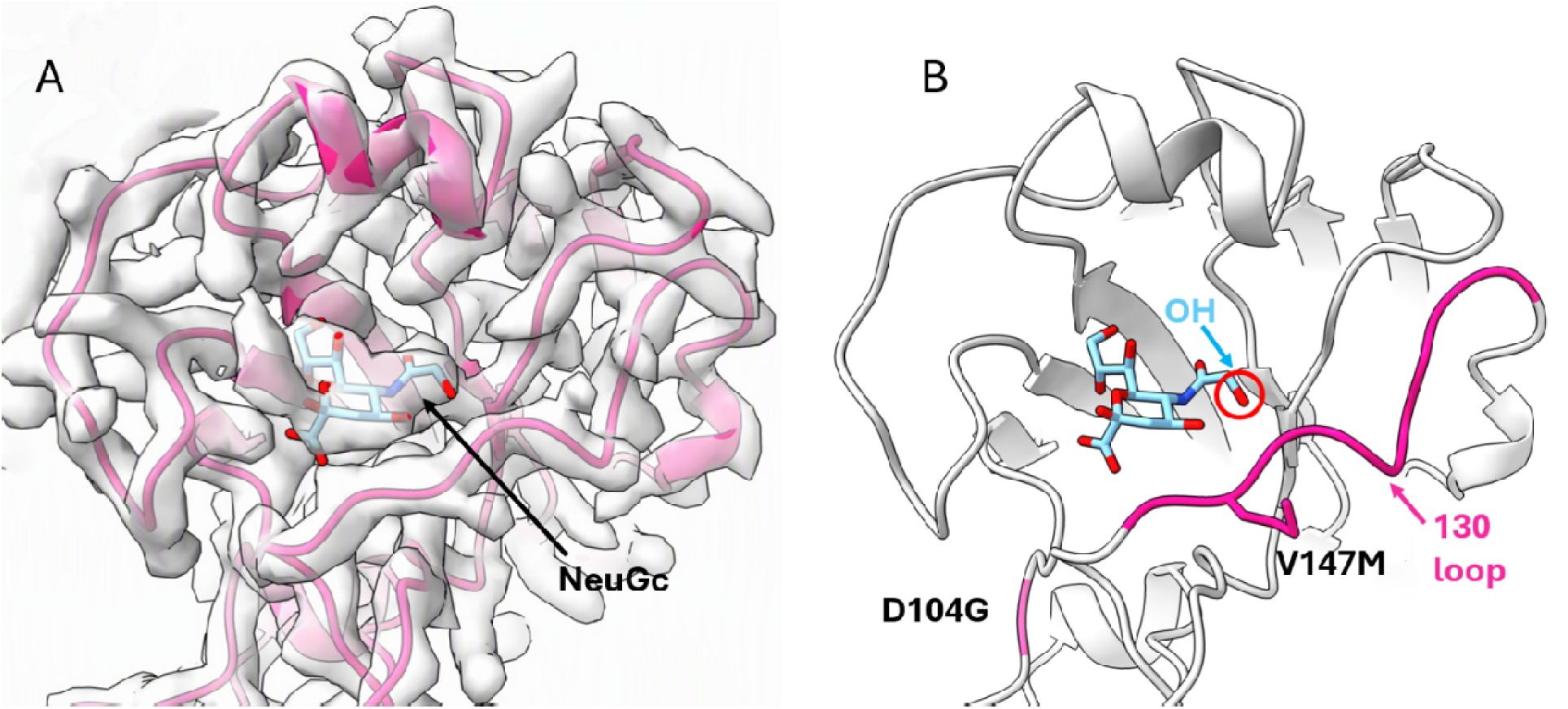
Structure of the cattle/California receptor binding domain bound to a NeuGc containing receptor analogue. **A)** Schematic representation within the cryo-EM density map. The ligand is coloured pale blue. B) Schematic representation where the 130 loop is coloured pink and key mutations are shown from Cattle/Texas.

## Discussion

Host-specific glycan chemistry beyond sialic acid linkage alone represents a major but often underappreciated determinant of influenza virus adaptation during cross-specie transmission. The ongoing H5N1 outbreak in dairy cattle provides a natural system to examine how these molecular features shape viral evolution during sustained mammalian transmission. Earl adaptation of H5N1 in cattle was driven primarily by rapid changes in the viral polymerase complex^22–24^. Here we show that continued evolution involves gradual selection and fixation of HA mutations that enable recognition of a sialic acid variant abundant in bovine tissues but not synthesised in birds and humans. Several of the most prevalent substitutions enhance replication in cattle tissues while being neutral or slightly attenuating replication in human airway models. In contrast to NeuGc-preferring viruses such as equine H7N7^17,18,26^, these mutations broaden receptor usage rather than switch specificity. This expanded receptor engagement may therefore preserve host range, consistent with spillover of H5N1 viruses carrying D104G and V147M into humans and poultry^33^.

We have also demonstrated that bovine respiratory tissues express abundant NeuGc-containing glycans, therefore may also be a site for ongoing adaptation which could facilitate more efficient respiratory replication and transmission, consistent with emerging reports of respiratory involvement in cattle infections^34^. Such changes could alter transmission dynamics within dair herds and complicate control efforts, while also increasing the risk of spread between cattle production systems. Several other influenza hosts, including horses, pigs and sheep, also express high levels of NeuGc. Although infections or serological evidence in these species during the current panzootic remain sporadic^35–37^, adaptation in one host may increase spillover potential into others. These dynamics highlight the importance of rapid control of livestock influenza outbreaks, as prolonged circulation may generate evolutionary changes that expand ecological opportunities for the virus.

Other selective pressures may also have contributed to the repeated emergence and fixation of these mutations. Both D104G and V147M have been associated with modest antigenic effects in H5N1^38^. Although these substitutions emerged when herds were likely immunologically naive, later fixation may in part reflect immune selection during reinfection as the outbreak progressed.

We further show that relevant bovine tissues contain substantial levels of α2–6-linked sialic acids. This observation aligns with recent reports that these tissues can support infection with both human and avian influenza viruses^39^, supporting the possibility that cattle could act as a mixing vessel for reassortment or adaptation of viruses with zoonotic potential. Although we did not observe mutations affecting α2–6 receptor engagement during this period, such adaptations warrant close monitoring as H5N1 continues to circulate in cattle.

Together, these findings suggest that host-specific modifications of sialic acids can act as powerful evolutionary filters shaping influenza receptor usage during mammalian emergence. Incorporating this broader glycan diversity into surveillance frameworks may therefore improve our ability to anticipate host shifts and assess zoonotic risk as influenza viruses continue to expand into new mammalian reservoirs.

## Materials and Methods

### Ethics and biosafety

Virus work was undertaken at either containment level 3 (CL3), SAPO4 (for whole H5N1 reverse genetics-derived viruses) or CL2 (for reverse genetics-derived viruses with HA and NA from H5N1, where the polybasic cleavage site of the H5HA was replaced with a monobasic cleavage site, and the remaining genes from the attenuated vaccine strain A/Puerto Rico/8/1934 (PR8)). Viruses carrying H5 HA with a multibasic cleavage site are categorised as specified animal pathogens order (SAPO) 4 and Advisory Committee on Dangerous Pathogens (ACDP) hazard group 3 by United Kingdom regulations. Work with these viruses was undertaken in a licensed CL3/SAPO4 facility of The Pirbright Institute under GMRA (BAG-RA-226). CL2 work with ACDP Hazard Group 2 recombinant influenza viruses was performed at the Roslin Institute under biological risk assessments BARA 1011 and GMRA 1811. All virus and GM risk assessments were approved by the appropriate internal committees, as well as the UK Health and Safety Executive (HSE) and, where necessary, the UK scientific advisory committee for genetic modification (SACGM).

Ethics for the use of the primary human airway epithelial cultures (hNECs) were as described previously^40^. Briefly, donors provided written consent and ethics approval was given through the Living Airway Biobank, administered through the UCL Great Ormond Street Institute of Child Health (REC reference: 19/NW/0171, IRAS project ID: 261511, Northwest Liverpool East Research Ethics Committee). Nasal brushings were obtained by trained clinicians from adult (30– 50 years) donors who reported no respiratory symptoms in the preceding 7 weeks. Brushings were taken from the inferior nasal concha zone using cytological brushes (Scientific Laboratory Supplies, CYT1050). All methods were performed following the relevant guidelines and regulations.

All the procedures involving embryonated eggs were undertaken in strict accordance with the guidance and regulations of the UK Home Office under project license number PP6471846. As part of this process, the work has undergone scrutiny and approval by the animal welfare ethical review board at The Pirbright Institute, incorporating the 3Rs and followed by ARRIVE (Animal Research: Reporting of in vivo experiment) guidelines for quality, reproducibility, and translatability of animal studies. Animal tissues were obtained from slaughtered dairy cows (Ethical approval URN 2024 2312-2). Generation of bovine explants came under approval from the University of Glasgow School of Biodiversity, One Health and Veterinary Medicine (EA26/25).

The deep mutational scanning experiments exclusively utilized pseudotpyed lentiviral particles (pseudoviruses) that can only undergo a single round of cell entry and so are not human pathogens. In addition, the deep mutational scanning identified mutations that increased binding to Neu5Gc receptors, which are expressed on bovine but not human cells; therefore these mutations would not increase the risk viruses pose to humans. Furthermore, mutations with this same phenotype (enhanced NeuGc binding) are already found widely in cattle-adapted H5N1 viruses.

### Bayesian phylogenetic analysis of HA

An H5N1 clade 2.3.4.4b dataset was compiled by querying the Global Initiative on Sharing All Influenza Data (GISAID; https://gisaid.org/)^41^ and NCBI Virus databases for all H5Nx clade 2.3.4.4b samples collected between November 1, 2021 – January 9, 2026, with available genomic sequences for all eight gene segments (accessed on January 9, 2026). Consensus genomes from raw reads deposited to the NCBI Sequence Read Archive and associated with BioProjects PRJNA1102327, PRJNA1122849, PRJNA1134696, PRJNA1219588, PRJNA1207547, PRJNA980729 were downloaded from the Andersen Lab’s avian-influenza github repository (https://github.com/andersen-lab/avian-influenza). Restriction to complete genomes enabled genotype assignment across all eight segments. The samples in each dataset were assigned a genotype using the GenoFLU genotyping tool^42^. Sequence datasets for genotype B3.13 were compiled for HA (n = 5,495). We further restricted the dataset to viruses sampled from dairy cattle and to those with complete collection dates, resulting in 3,929 HA sequences.

We used BEAST X v10.5.0^43^ to perform phylogenetic inference of the HA segment. We used a non-parametric skygrid prior^44^ with 23 gridpoints and a cutoff of 2.0 years, a strict clock with an informative normal prior with a mean of 4.8×10^−3^ substitutions per site per year and a standard deviation of 3.11×10^−4^ substitutions per site per year, a codon-partitioned model with a GTR substitution model for each codon position, and Hamiltonian Monte Carlo gradient-based sampling for node ages^45^. We ran three chains of 100 million generations, subsampling every 10 thousand iterations to continuous parameter log and tree files. Convergence and mixing were assessed in Tracer v1.7.2^46^ and 10% of each chain was discarded as burn-in. The three chains were then combined and downsampled using LogCombiner, to construct an empirical tree distribution with a total of 509 trees. All relevant ESS values were >150.

We next used the robust counting framework^47^ in BEAST X to infer the number of substitutions on branches and their neutral expectations. We used the empirical tree distribution described above, with the remaining parameterizations the same. We simulated one MCMC chain of one million generations, subsampling every one hundred iterations to continuous parameter log and tree files. Convergence and mixing were assessed in Tracer and the first 10% of the chain was discarded as burn-in. All relevant ESS values >150.

All phylogenetic trees were processed using TreeSwift v1.1.42^48^ and visualized using baltic v0.2.2^49^.

Median dN/dS values from the robust counting analysis were mapped onto a high-resolution protein structure for HA (PDB: 9DWE^50^) using PyMOL v3.1.0.

### Tissues and Glycomics

Cattle tissues were collected under ethical approval URN 2024 2312-2. The following samples were collected from a Holstein-Friesian cow: mammary tissue surrounding the gland cisterns; internal layer of the teat; internal section of trachea; pieces of lung lobes. Tissues were washed with PBS and stored at −80°C until required. Mammary and teat tissues from 4 cows, and trachea and lung tissues from 2 cows were analysed. Both lactating cattle (animal 1 and 3) and non-lactating cattle (2 and 4) were analysed.

Approximately 1 g (wet weight) of the tissues were washed multiple times with ice cold PBS, then processed according to the protocol by Jang Lee at al., 2006^51^. Briefly, the tissue samples were sonicated to lyse the cells, then samples were dialysed and lyophilised. Reduction and carboxymethylation reactions were carried out on the samples, which were then treated with trypsin (Sigma) and the resultant glycopeptides were purified by Sep-Pak C18 chromatography. PNGase-F (Roche) was used to release the N-Glycans which were subsequently purified by Sep-Pak C18 chromatography. O-glycans were released by a reductive elimination reaction with potassium borohydride (KBH4) and potassium hydroxide (KOH) then subsequently purified using a desalting column of Dowex beads as well as Sep-Pak C18 chromatography. Both N- and O-glycans were permethylated prior to analysis. All MS and MS/MS data were acquired on a Bruker neofleX MALDI-TOF/TOF instrument equipped with 10 kHz smartbeam 3D laser and controlled by flexControl 5.0 software. Spectra of permethylated glycans were acquired in positive reflector mode. N-glycan spectra were accumulated by adding 12,000 laser shots and setting the m/z detection range to 1000 – 10,000. O-glycan spectra were accumulated by adding 12,000 laser shots and setting the m/z detection range to 300 – 3,000. External m/z calibration was performed using a permethylated dextrose ladder. MS/MS spectra of selected glycan molecular ions were acquired in positive reflector mode. MS and MS/MS spectra were processed in flexAnalysis 5.0 software (peak finding and baseline correction). [M+Na]^+^ molecular ions were annotated using a predefined database of possible structures and with the assistance of GlycoWorkbench^52^ (raw data in Supplementary tables 3-5). Annotations were based on glycan composition and knowledge of biosynthetic pathways. MS/MS analysis was used to confirm predicted assignments (Supplementary Figure S4) and Adobe Illustrator was used to prepare the figures. For each tissue, the glycans were treated with Sialidase-S or Sialidase-A (Agilent Technologies) to determine the sialic acid linkage configurations. These samples were then permethylated prior to analysis. The proportions of NeuAc and NeuGc, and their respective linkage configurations were calculated using R. The relative intensity of each glycan was normalised to the sum of the relative intensities of all the glycans in the sample. The normalised intensities were then weighted based on the number of NeuAc and NeuGc residues present on each sialylated glycan. The data from samples digested with Sialidase-S, up to the highest m/z glycan detected in the untreated sample, was used to determine the proportions of α2-3 and α2-6 linked sialic acids: the weighted normalised intensities of sialylated molecular ion peaks remaining after sialidase-S digest were summed to give the proportion of α2-6 linked NeuAc and NeuGc. The weighted normalised intensities of sialylated molecular ions in the sialidase-S sample were subtracted from the weighted normalised intensities of sialylated molecular ions from the untreated sample in order to calculate the proportion of α2-3 linked NeuAc and NeuGc.

### Cells, organ cultures and eggs

Human Embryonic Kidney 293T (293T) and Madin-Darby Canine Kidney (MDCK) were maintained in Dulbecco’s modified Eagle medium (DMEM) supplemented with 10% fetal bovine serum (FBS), 1% sodium pyruvate, and 1% Pen-Strep. Human lung adenocarcinoma (Calu-3) cells were maintained in DMEM with 10% FCS, 1% Pen-Strep, 2 mM L-glutamine. All cells were maintained at 37°C and 5% CO2.

293T-CMAH and 293T-GFP control cells were generated as previously described^30^. Briefly, 293T cells were transduced with a lentiviral construct, RRL.sin.cPPT.SFFV/ptCMAH.IRES-puro.WPRE or RRL.sin.cPPT.SFFV/GFP.IRES-puro.WPRE^53^, expressing codon optimised (human) chimpanzee CMAH (NP_001009041.1) or GFP, and selected using puromycin.

HEK293 ΔST3GAL1/2/3/4/5/6, ΔST61/2 cells were a kind gift from Professors Henrik Clausen and Yoshiki Narimatsu of the University of Copenhagen^31^.

Specific pathogen free embryonated hen’s eggs were purchased from Valo. Eggs were matured to 10-days-old and checked for viability before use for propagating viruses.

Primary human nasal epithelial cultures (hNECs) were generated as described previously^40^. Briefly, basal epithelial cells from nasal brushings were expanded on mitotically inactivated 3T3-J2 fibroblasts to passage 1 and cryopreserved. Cells were thawed as required, expanded with 3T3-J2 fibroblasts to passage 2, and seeded onto collagen I–coated, semi-permeable membrane supports (3 × 10^5 cells/6.5 mm, 0.4 µm pore size, Transwell; Corning). Differentiation was performed under air–liquid interface conditions in PneumaCult™-ALI medium for 4 weeks.

Bovine mammary explants were produced as previously described^22,23^. Briefly, Tissue was collected from Sandyford Abattoir from udders of commercially slaughtered cows free from mastitis or any antibiotic treatment. Excess tissue was removed, teat and gland cisterns were dissected and transferred to transport media comprising of chilled DMEM supplemented with 100 μg/ml Penicillin/ 100 μg/ml streptomycin and 5 μg/ml fungizone. Every 45 minutes, tissues were transferred to 500ml of fresh transport media. Inside a class II biological safety cabinet, explants were made using a 5mm biopsy punch. Biopsies were added to 24 well plates with 500 ul of transport media supplemented with 10% FBS and incubated for 24h at 37°C, 5% CO2 before infection.

### Plasmids

Reverse genetics plasmids of the viruses used in this study were produced as previously described (Table 1)^6,23,53^. Mutants were generated by site directed mutagenesis.

**Table 1.** Virus strains used in this study.

| Virus strain name | Alias | Accession number(s) | Whole virus, 2:6 (PBCS) or pseudovirus |
| --- | --- | --- | --- |
| A/dairy_cow/Texas/24_009108-001/2024(H5N1) | cattle/Texas | EPI_ISL_19094762<sup>4</sup> | Full virus, 2:6 MBCS and pseudovirus |
| A/cattle/CA/24-036570-001-original/2024(H5N1) | cattle/California | XLZ31360.1 | Full virus (HA only), Monobasic H5HA, and pseudovirus |
| A/equine/Detroit/3/19<br>64(H7N7) | eqH7N7 | AGT18811.1,<br>AGT18813.1 | Pseudovirus only |
| A/England/195/2009(<br>H1N1) | Eng195 | EPI_ISL_29994 | Pseudovirus only |

Plasmids used for pseudovirus assays were used as previously described^24,54^. HA and NA plasmids for cattle/Texas, equine H7N7 and chimpanzee CMAH (NP_001009041.1) were human codon optimised and synthesised by Genscript in a pcDNA3.1 background.

Reverse genetics plasmids for the H5HA of cattle/Texas with the furin cleavage site replaced with a monobasic cleavage site, and NA were used as previously described^6,53^. Mutants of cattle/Texas HA, and the cattle/California HA were generated by site directed mutagenesis. H5HA (again with a monobasic cleavage site replacing the polybasic cleavage site) and N1NA of AIV07 used as previously described^6^.

### Biophysical receptor binding assays

Viruses were purified as previously described^6^. Briefly, 400 ml of clarified, virus containing allantoic fluid harvested from embryonated chicken eggs was concentrated by ultracentrifugation at 27,000 rpm for 2 hours at 4 °C. Pelleted virus was resuspended and homogenised using a glass homogeniser, then passed through a 30-60% sucrose gradient by ultracentrifugation at 27,000 rpm for 2 hours at 4 °C. Virus containing fractions were harvested, and diluted in PBS, and ultracentrifuged at 27,000 rpm for 2 hours at 4 °C to remove sucrose. Finally, the virus-containing pellet was resuspended in PBS. Enzyme-linked immunosorbent assay (ELISA) was used to quantify the amount of virus comparing NP to a known standard^55^.

Biotinylated polymeric probes containing α-2-3-sialyllactosamine (3SLN), α2-6-sialyllactosamine (6SLN), Neu5Gcα3′LN (3S[Gc]LN) Neu5Gcα6′LN (6S[Gc]LN) were purchased from GlycoNZ. Receptor binding assays using bio-layer interferometry were performed as previously described^6^. Briefly 100 pmol of purified virus was diluted with Hepes buffered saline with EDTA and tween 20 (HBS-EP; (TEKnova) containing 10 μM oseltamivir carboxylate (Roche) and 10 μM zanamivir (GSK). Receptor binding kinetics were measured using the Octet® R8 system (Sartorius) and streptavidin biosensors (Sartorius). Virus binding affinity was normalised to fractional saturation plotted against sugar loading values^56,57^.

### Antisera preparation

Cattle/Texas WT virus was propagated in in 9-day old chicken embryos, the virus was then harvested and inactivated with β-propiolactone (BPL) and concentrated by ultracentrifuge at 27,000 rpm for 2 hours. Three-day-old specific pathogen free (SPF) chickens were vaccinated with 1024 haemagglutinating unit (HAU) inactivated virus mixed with oil emersion adjuvant (Montanide; Seppic) at a 7:3 ratio (adjuvant : virus). The chickens were boosted with the same amount of antigen 10 days after. The antisera were then harvested 42 days post-vaccination for subsequent glycan array detection.

### Glycan arrays

Commercial chemicals and solvents were of the highest grade available. All chemicals were from Merck unless otherwise stated. Thin layer chromatography (TLC) was performed on Silica Gel F254 (Merck) with detection by UV-light. High performance TLC (HPTLC) for glycolipids (GLs) and neoglycolipids (NGLs) was performed on Silica Gel plates (Merck, Dorset, UK). For HPTLC analyses, NGLs were applied onto the silica gel plates with a TLC applicator (Camag Linomat 5, Muttenz, Switzerland) and the TLC tank was equilibrated with the developing solvents (as indicated) for 30 min prior to chromatography. Lipids were detected by UV-light after staining with primulin solution1 and glycans were visualized by staining with orcinol solution1. The quantitation based on readouts from the chromatogram plates was performed using a Camag TLC Scanner 3. HPLC purification of derivatized glycans was performed on a WATERS E2695 system with 2489 UV detector. For normal phase (NP) HPLC, an Amide HPLC column (3.5 μm, 4.6 mm x 250 mm, Xbridge, WATERS) was used. MS spectra were recorded with a Shimadzu 8030 MALDI instrument.

For conversion to NGLs (Supplementary Figure S18), 10 nmol of dry azido glycans were each dissolved in 40 μL CHCl3:MeOH:100 mM ammonium acetate in H2O 25:25:8 containing 40 nmol of Dibenzocyclooctyne–1,2-dihexadecanoyl-sn-glycero-3-phosphoethanolamine (DBCO-DHPE) (REF FAA). The reaction was kept at R.T. for 16 h and the NGLs purified by NP HPLC with an amide column using CHCl3:MeOH:H2O (Solvent A 130:70:9 to Solvent B 10:20:8). A short gradient was used here to effectively remove excess DBCO-DHPE. The column was then kept in 95% solvent A for 5 min to elute DBCO-DHPE followed by switching to 90% B for 4 min isocratically to elute the NGL product. The products were quantified on HPTLC (Supplementary Figure S19) following a previously published method^58^. The corresponding MS spectra are presented in Supplementary Figure S20.

The noncovalent array was prepared in parallel using the NGL form of 12 NGL probes. The list of 6 GBPs investigated and assay conditions are in Supplementary Table 6. Details on the arraying instrument, printing conditions, array layout, imaging and data analysis are in the MIRAGE document (Supplementary Table 7). The source data are in the Supplementary Tables 6 and 7. The pre-wetted slide was blocked in HEPES-buffered saline (HBS) containing 2% bovine serum albumin (BSA) for 1 h. The slide was then washed with HBS and incubated with 100 pmol of purified virus supplemented with 100 µM oseltamivir and 100 µM zanamivir for 90 min. Afterward, the slide was washed four times with HBS, fixed with 4% paraformaldehyde (PFA) for 30 min, and incubated with chicken polysera against cattle/Texas WT (diluted 1:100) for 1 hour. The slide was then incubated with biotinylated anti-chicken IgY (1:200; BA-9010-1.5, Vector Laboratories) for 1 hour. Finally, the slide was washed three times with HBS and incubated with streptavidin-AF647 (1:1000; S21374, ThermoFisher Scientific) for 30 min.

### Pseudovirus entry assays

Pseudovirus were produced as previous described^59^. Briefly, HEK 293T cells seeded in 6-well dishes co-transfected using lipofectamine 3000 (Invitrogen) at ∼70% confluency with 0.6 µg of the luciferase reporter constructs (pCSFLW), 0.4 µg of the HIV packaging plasmid pGAG-POL, and either 0.4 µg of VSV-G (VSV-G only) or 0.4 µg of the defined HA, 0.2 µg of TMPRSS11D (non-H5 only, e.g. H1N1 and H7N7) and 0.1 µg of matched NA. Pseudovirus-containing supernatants were collected and pooled at 48 and 72 hours post-transfection, clarified by centrifugation, aliquoted and frozen at –80°C.

Entry assays into 293T-CMAH vs 293T-GFP cells was performed by overlaying 100 µL cells at a concentration of 10^5^ cells/ml onto pseudovirus diluted in oseltamivir-containing media (final concentration 0.5 µM). Cells were then left for 48 hours at 37°C, 5% CO_2_ then lysed and read using Brightglo reagent (Promega) and a Glomax discover plate reader (Promega). Data for each pseudovirus was normalised to the GFP control.

Siaylyl-transferase reconstitute assays were adapted from the previously described protocol^59^. 293 cells lacking endogenous sialyl-transferase expression (293 ΔST3GAL3/4/6 ΔST61/2^30^) were transiently transfected with 100 ng of expression plasmids encoding either empty vector, human ST3GAL4, ST6GAL1, mixed with 400 ng of either empty vector or chimpanzee CMAH. 24 hours later cells were scraped, diluted to 10^5^ cells/ml, transferred into 96 well plates (100 µL/ well) and mixed with 100 µL of the named pseudoviruses diluted in oseltamivir (final concentration 0.5 µM). Cells were then left for 48 hours at 37°C, 5% CO_2_ then lysed and read using Brightglo reagent (Promega) and a Glomax discover plate reader (Promega). Data for each pseudovirus was normalised to the empty vector control.

### Virus replication kinetics

Infection of explants was performed as follows: individual explants were transferred to 96-well plates and infected with 5000 PFU of virus in 100 µl of serum free DMEM for 2 hours at 37 °C. Explants were transferred to clean 24-well plates, washed 3x with warm PBS and overlayed with 1 ml of DMEM supplemented with 0.14% fraction V BSA and 1 µg/ml TPCK trypsin. Time points were collected at the indicated times post-infection.

Confluent monolayers Calu-3 cells were washed once with PBS and infected at multiplicity of infection (MOI) 0.01 with virus diluted in serum-free medium for 1 hour at 37 °C. Inoculum was replaced with serum-free media. Supernatants were collected at various times post infection and stored at −80°C until infectious titres were determined by plaque assay.

Infection of primary human airway cultures was performed as follows. Before infection cells were washed with Dulbecco’s phosphate-buffered saline supplemented with calcium and magnesium (DPBS++) to remove mucus and debris. Cells were infected with 200 µl of virus in DPBS++ at an MOI of 0.01 and incubated at 37°C for 1 h. Inoculum was removed, and cells were washed twice with DPBS++. Time points were taken by adding 200 µl of DPBS++ and incubating for 10 mins and 37°C before removal and titration. The time course was performed at 37 °C, 5% CO2. Time points were taken at 24-, 48-, 72-hours post-infection. Infectious titres were determined by plaque assay on MDCKs.

### Pseudovirus deep mutational scanning

Generation of pseudovirus-based H5 HA deep mutational scanning library was described previously^24^. This library uses pseudotyped lentiviral particles that can only undergo a single round of cell entry, and so are not human pathogens and therefore provide a way to study H5 HA mutations at biosafety-level 2. Deep mutational scanning library was made in the genetic background of the HA of A/American Wigeon/South Carolina/USDA-000345-001/2021(H5N1) strain. Cell entry effect measurements were performed as described previously^24^. In brief, 3 million transcription units of HA pseudotyped libraries and 10 million transcription units of same libraries pseudotyped with VSV-G were used to infect HEK-293 and chimpanzee CMAH (293-CMAH) overexpressing cell lines. 15 hours post infection cells were collected, non-integrated pseudovirus genomes were recovered and Illumina sequencing libraries were prepared, as described previously^24^. Two biological library replicates were used in these experiments.

Effects of mutations on cell entry were calculated using log enrichment ration: 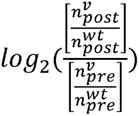, where 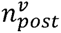 is variant *n* count post-infection (HA pseudotyped library infection) 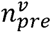 is variant *n* count pre-infection (VSV-G pseudotyped library infection) and 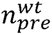 and 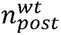 are counts for unmutated HA variants pre and post-infection, respectively. The multidms package^60^ was used to fit global epistasis model to variant cell entry scores and estimate mutation-level effects on cell entry. Generation of 293-CMAH cells used in these experiments was described previously^29^. The GitHub repository with computational pipeline used for analysis is at https://github.com/dms-vep/Flu_H5_American-Wigeon_South-Carolina_2021-H5N1_DMS_NeuGc and numerical values for cell entry effects are at https://github.com/dms-vep/Flu_H5_American-Wigeon_South-Carolina_2021-H5N1_DMS_NeuGc/blob/master/results/summaries/entry_in_NeuAc_vs_NeuGc_cells.csv..

### Expression and purification of HA ectodomains

Haemagglutinin ectodomains were expressed in Expi293 cells (amino acids 1-502) with a C-terminus TEV_foldon_twinstrep tag. Codon optimised DNA was synthesised by Genscript into a pcDNA3.4 vector. Six days after transfection the HA trimers were captured from the cell culture supernatant with Strep-Tactin XT resin (IBA Life Sciences) and subsequent elution (using the manufacturers protocol) was followed by size exclusion chromatography on a Superose 6 increase column (Cytiva) in 25mM Tris pH 8, 150 mM NaCl and 0.5 mM TCEP.

Samples were then incubated overnight at 37°C with Sialidase A (Agilent Technologies) at pH 6 (buffer provided) to remove autoglycan occupation of the sugar binding site. This was followed by centrifugation for 10 minutes at 20,000 G to remove aggregated protein. Size exclusion chromatography on a Superose 6 increase column (Cytiva) was then repeated in 25 mM Tris pH 8, 150 mM NaCl with 0.5 mM TCEP.

### CryoEM

NeuGc occupied samples were incubated with 2 mM 3SLN(NeuGc)^61^ for 10 minutes at room temperature, and 0.1% Octyl Glucoside was used to supplement all samples. Plunge freezing onto homemade graphene oxide grids (GO layer on R2/2 Cu 300 mesh (Quantifoil)) was performed on a Leica GP2 (Leica microsystems) with auto-blotting plus 1mm, a wait time of 20s, and 6s blot time. 4 μl of each sample was blotted at 20 μg/ml in 100 % humidity and 4°C.

A 300 kV Titan Krios transmission electron microscope fitted with a Falcon 4 detector (Thermo Fisher) and an energy filter was used for data collection in counting mode using EPU. The total dose was 41.5 e-/Å2 dose fractionated into 31 frames. The defocus range was from −1.5 μm to −3.5 μm.

Data sets were processed on Relion 5.0^62^ and cryoSPARC v.4.7.1^63^ and model building was done using Coot1^64^ using both PDB 9DIP^65^ and Alphafold2^66^ de novo models as references. Models are currently being refined using Phenix and validated using Molprobability for submission to the PDB. Figures were generated using UCSF Chimera X^67^. Cryo EM structure of hemagglutinin from the Influenza Virus A/cattle/CA/24-036570-001-original/2024(H5N1) with bound NeuGc can be found under PDBID: 34WP.

## Acknowledgements and funding

The authors would like to thank Professors Henrik Clausen and Yoshiki Narimatsu of the University of Copenhagen for kindly sharing the HEK293 ΔST3GAL1/2/3/4/5/6, ΔST61/2 cells. The authors want to thank Professor John B. Lowe, of the university of Michigan Medical school for kindly gifting murine E-Selectin. The authors would also like to thank Professor Colin Parrish for advice, input and reagent sharing during the drafting of this paper.

We gratefully acknowledge all data contributors, i.e., the Authors and their Originating laboratories responsible for obtaining the specimens, and their Submitting laboratories for generating the gene sequence and metadata and sharing via the GISAID Initiative, as well as NCBI and SRA on which this research is based.

J.A.H, J.Y., D.H. J.Y., K.R.V., J.L.Q., L.A., H. J. K., S.R., J.J., I.H.B., D.H.G., Y.L, M.I., W.S. B., S.M.H. and T.P.P were supported by the UK Medical Research Council/Department for Environment, Food and Rural Affairs (Defra, UK) FluTrailMap-One Health consortium [MR/Y03368X/1]. J.A.H., J.Y., K.R.V., J.L.Q., J.J., A.N., I.H.B., M.I., W.S.B, S.M.H., and T.P.P. are also supported by the Biotechnology and Biological Sciences Research Council (BBSRC)/DEFRA ‘FluTrailMap’ consortium [BB/Y007298/1]. J.Y. J.L.Q., J.J., I.H.B., M.I., W.S.B., and T.P P. were previously supported by the BBSRC/Defra “FluMAP” consortium (BB/X006123/1; BB/X006166/1). J.Y., J-R.S., K.R.V., S.J., D.D., S.R., I.H.B., M.I. and T.P.P. were funded by the BBSRC via the Pirbright Institute’s Strategic Programme Grants (ISPGs) [BBS/E/PI/230002A; BBS/E/PI/230002B], and BBSRC National Bioscience Research Infrastructure: High Containment and Low Containment Services and Science Platforms grants [BBS/E/PI/23NB0004, BBS/E/PI/23NB0003]. N.H., A.N. and J.J. were also supported by the Defra and the Devolved Administrations of Scotland and Wales, through SE2227 ‘FluFocus’. T.P, S.J. and D.D., received support from the Royal Society research grant (RGS\R2\242118) and Houghton Trust small project research grant (HT/SPRG/23/04). J.Y and Y.L acknowledge funding from The UK Medical Research Council grant [MR/R010757/1]. J.D.B. is an investigator with the Howard Hughes Medical Institute. The Bruker neofleX MALDI-TOF/TOF instrument was funded by the BBSRC (UKRI2253). P.L. acknowledges support by the Research Foundation - Flanders (‘Fonds voor Wetenschappelijk Onderzoek - Vlaanderen’, G005323N, G051322N and G010326N). PRM was funded by MRC (MC_UU_0034/2, and MC_UU_0034/3) and BBSRC (BB/V004697/1). B.R.W. supported under a Centers of Excellence for Influenza Research and Response (CEIRR) grant 75N93021C00045 to Colin R. Parrish and Brian R. Wasik. J.E.P. acknowledges funding from The Rockefeller Foundation (PC-2022-POP-005). RP and PD acknowledge Institute Strategic Programme grant funding from the BBSRC (BBS/E/RL/230002A and BBS/E/RL/230002A). They also acknowledge co-funding by the European Union under project 101136346 EUPAHW

Views and opinions expressed are however those of the author(s) only and do not necessarily reflect those of the European Union or the European Research Executive Agency. Neither the European Union nor the granting authority can be held responsible for them.

