## Supplementary Figures for "Bovine H5N1 influenza viruses have adapted to more efficiently use receptors abundant in cattle"

**Supplementary Figures and Tables**

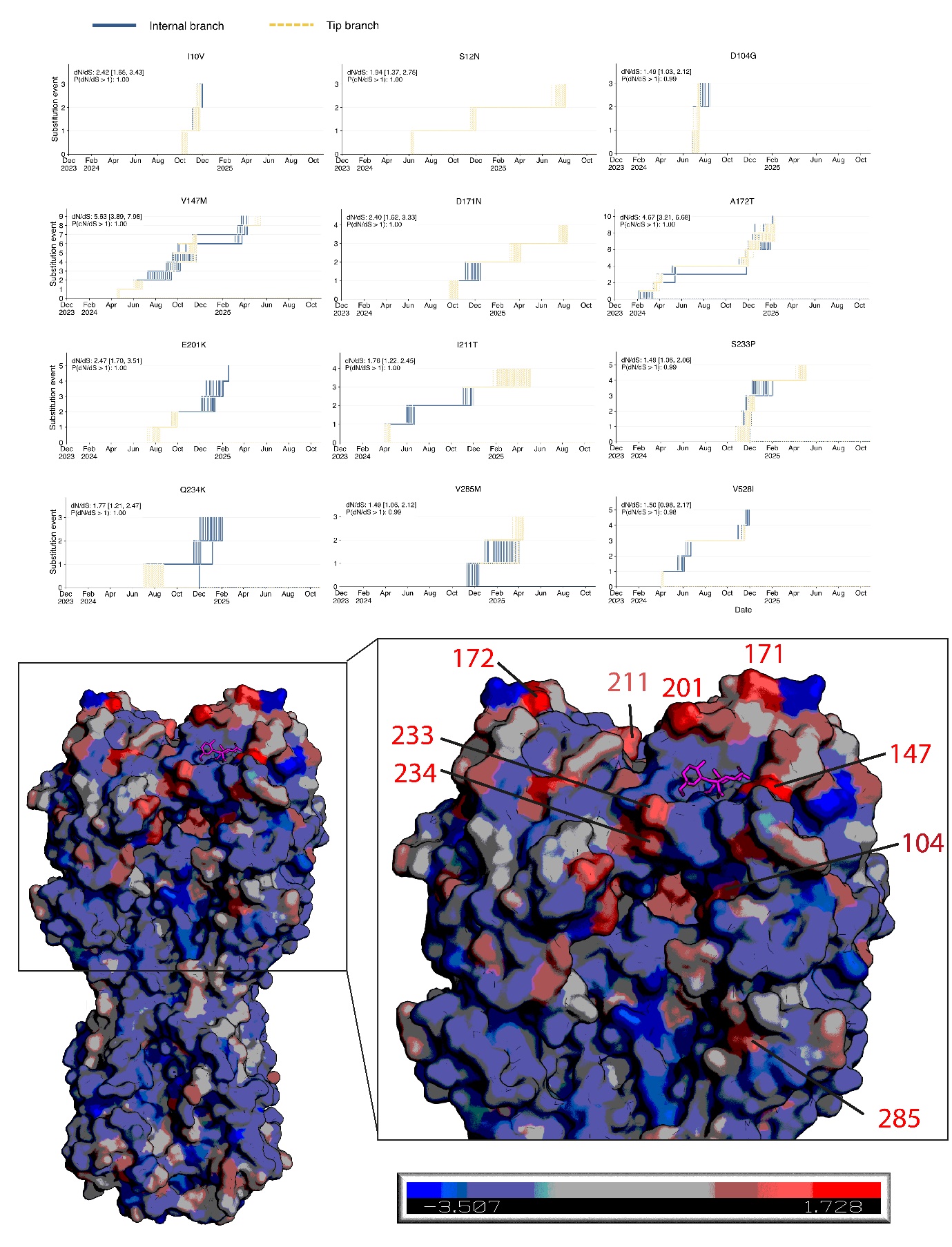

**Supplementary Figure S1. Haemagglutinin sites with signatures of positive selection.** Top panels **-** Each panel showcases the repeated emergence of an amino acid substitution across internal branches (solid dark blue lines) and tip branches (dotted yellow lines). Each stepwise path corresponds to the evolutionary trajectory of these substitutions in one posterior sample (n=509). The *dN/dS* value (median and 95% highest posterior density [HPD]) and its probability of being above 1 for the given substitution are indicated at the top of each panel. Only shown are amino acid substitutions inferred to have occurred at least twice and with high posterior support (>0.975) of *dN/dS* being above 1. **Bottom Panel –** sites under positive selection mapped on to the HA structure (PDB: 9DWE) in red, the alpha2,6-linked sialylated receptor analogue, LSTa, shown in purple.

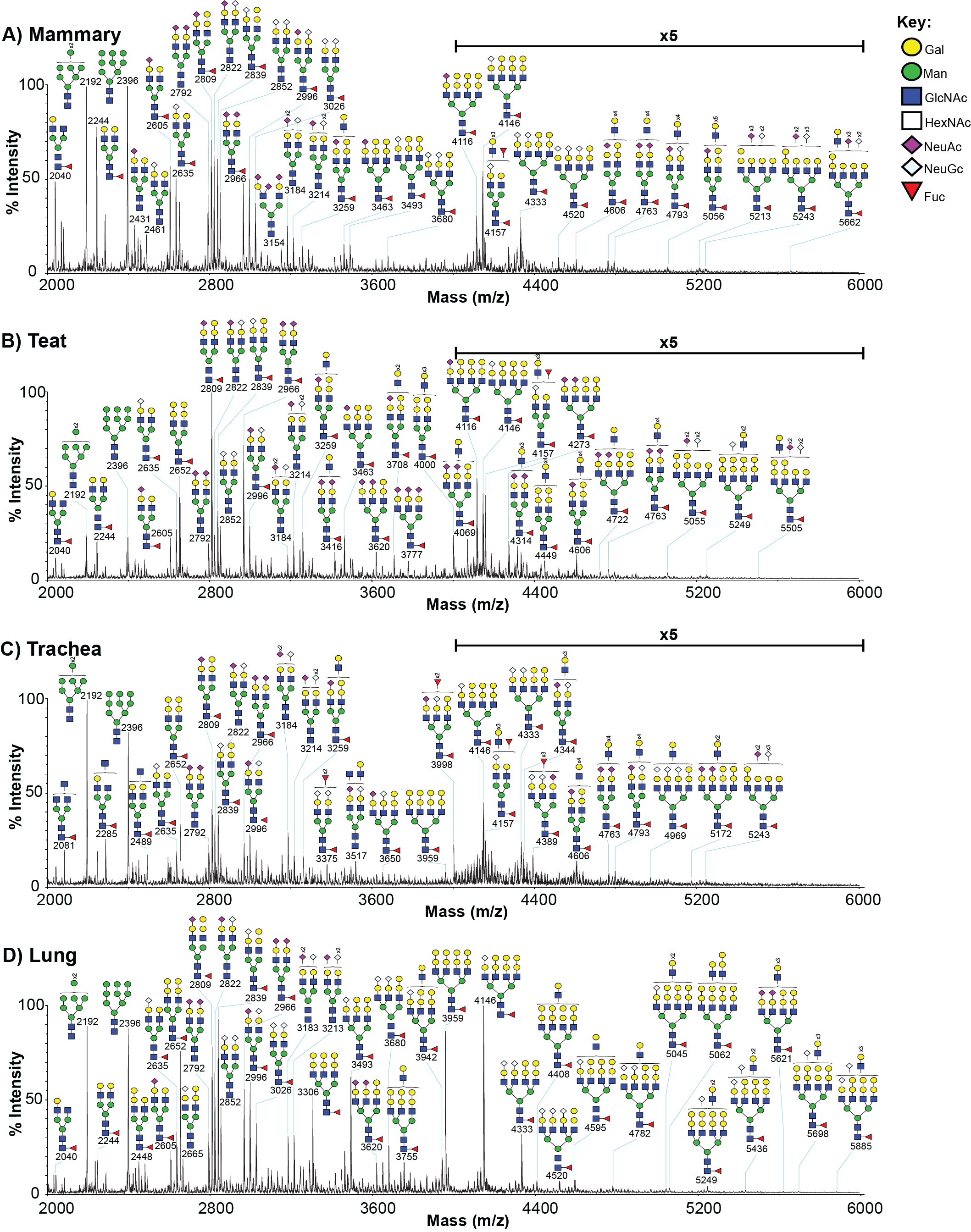

**Supplementary Figure S2: Annotated MALDI-TOF spectra of N-glycans from a second cow.**

Annotated MALDI-TOF spectra of bovine **A.** mammary, **B.** teat, **C.** trachea and **D.** lung tissue N-glycans from a second, non-lactating cow. Annotations show [M+Na]^+^ molecular ions. Peak annotation is based on composition, biosynthetic knowledge and MS/MS analysis.

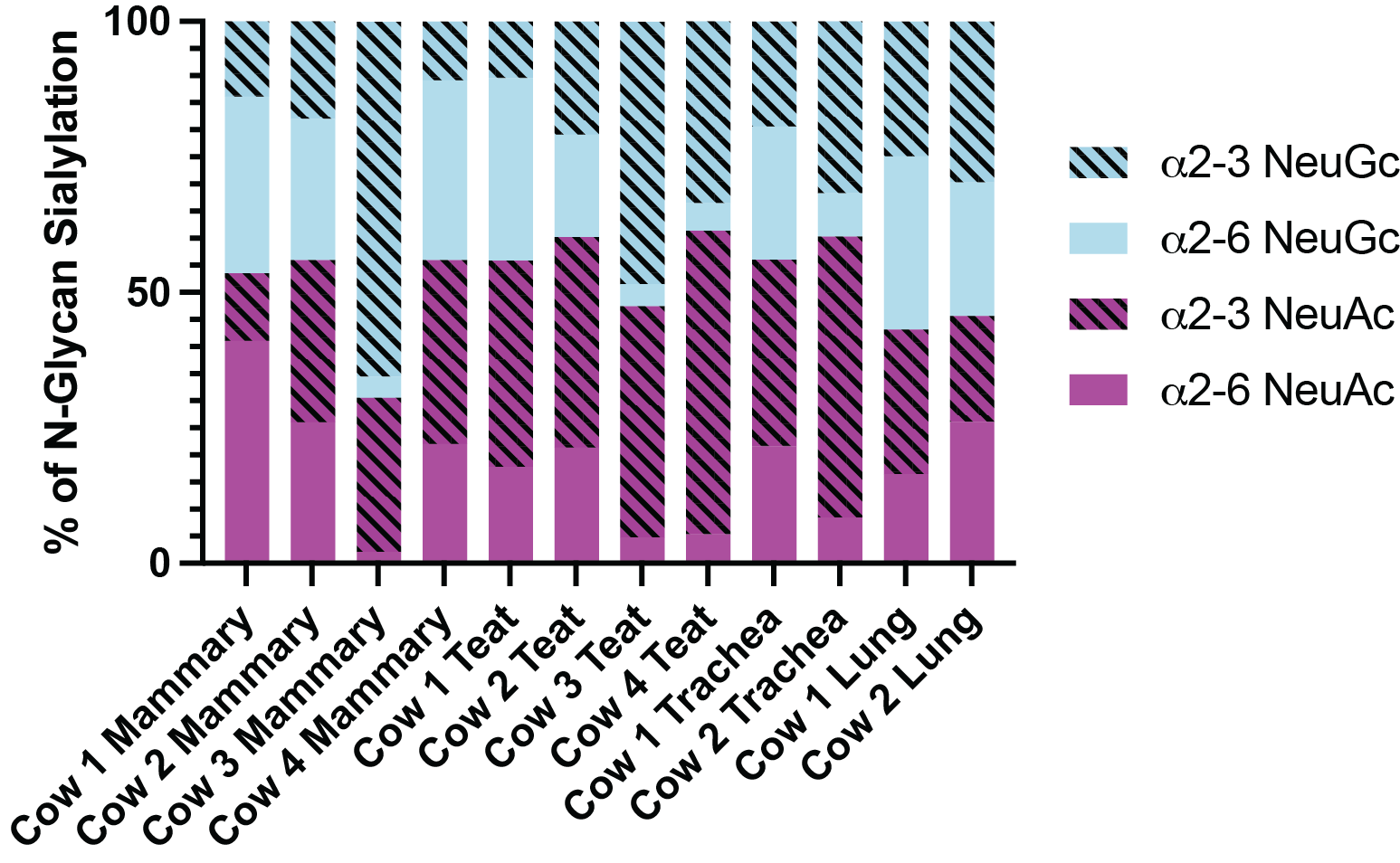

**Supplementary Figure S3: Proportions of α2-3 and α2-6 linked sialic acid in bovine tissue N-glycans.**

Summarised data is from lactating (cows 1 and 3) and non-lactating cows (cows 2 and 4).

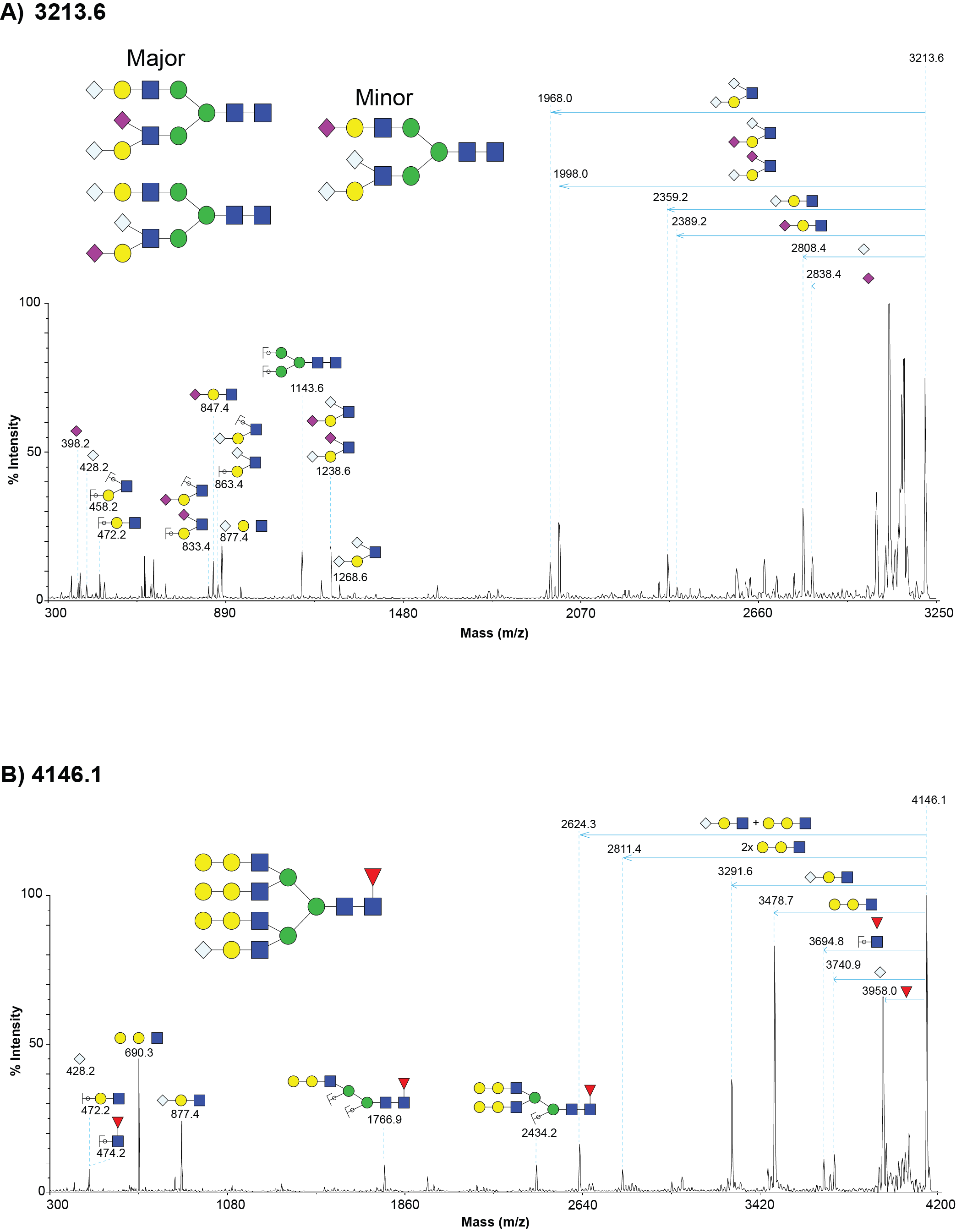

**Supplementary Figure S4: MALDI TOF/TOF MS/MS spectra of cow mammary N-glycans.**

MALDI TOF/TOF MS/MS spectra of A) *m/z* 3214 from B) *m/z* 4146 from lactating cow mammary tissue (cow 1) showing example MSMS data of a sialylated N-glycans. Annotations show [M + Na]^+^ fragment ions. Horizontal arrows correspond to the loss of the indicated fragment from the molecular ion.

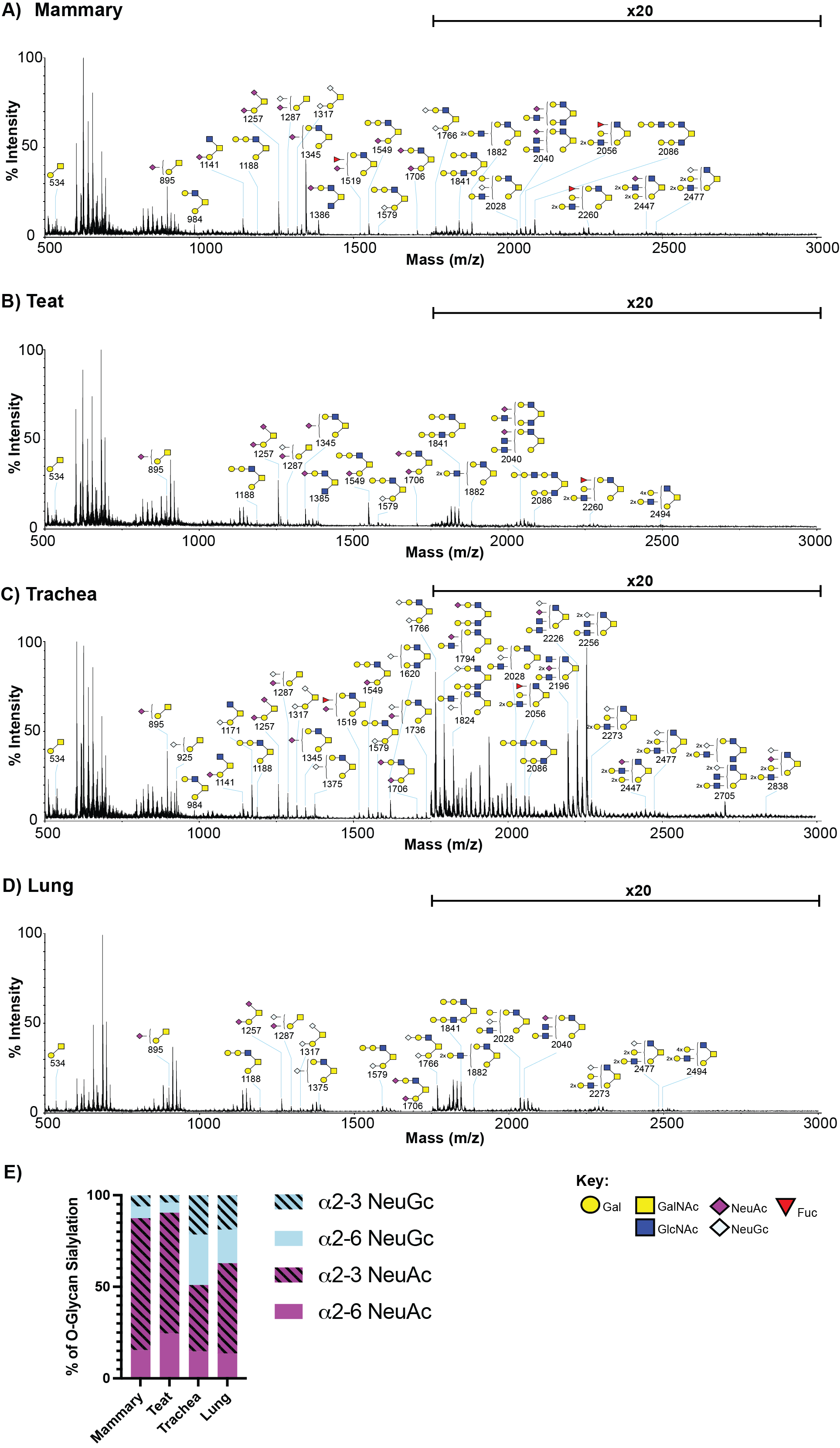

**Supplementary Figure S5: Annotated MALDI-TOF spectra of bovine tissue O-glycans.**

A) Mammary, B) teat, C) trachea and D) lung tissue O-glycans from a non-lactating cow (cow 2). Annotations show [M+Na]^+^ molecular ions. Peak annotation is based on composition, biosynthetic knowledge and MS/MS analysis. E) Calculated proportions of NeuAc and NeuGc in bovine O-glycan samples.

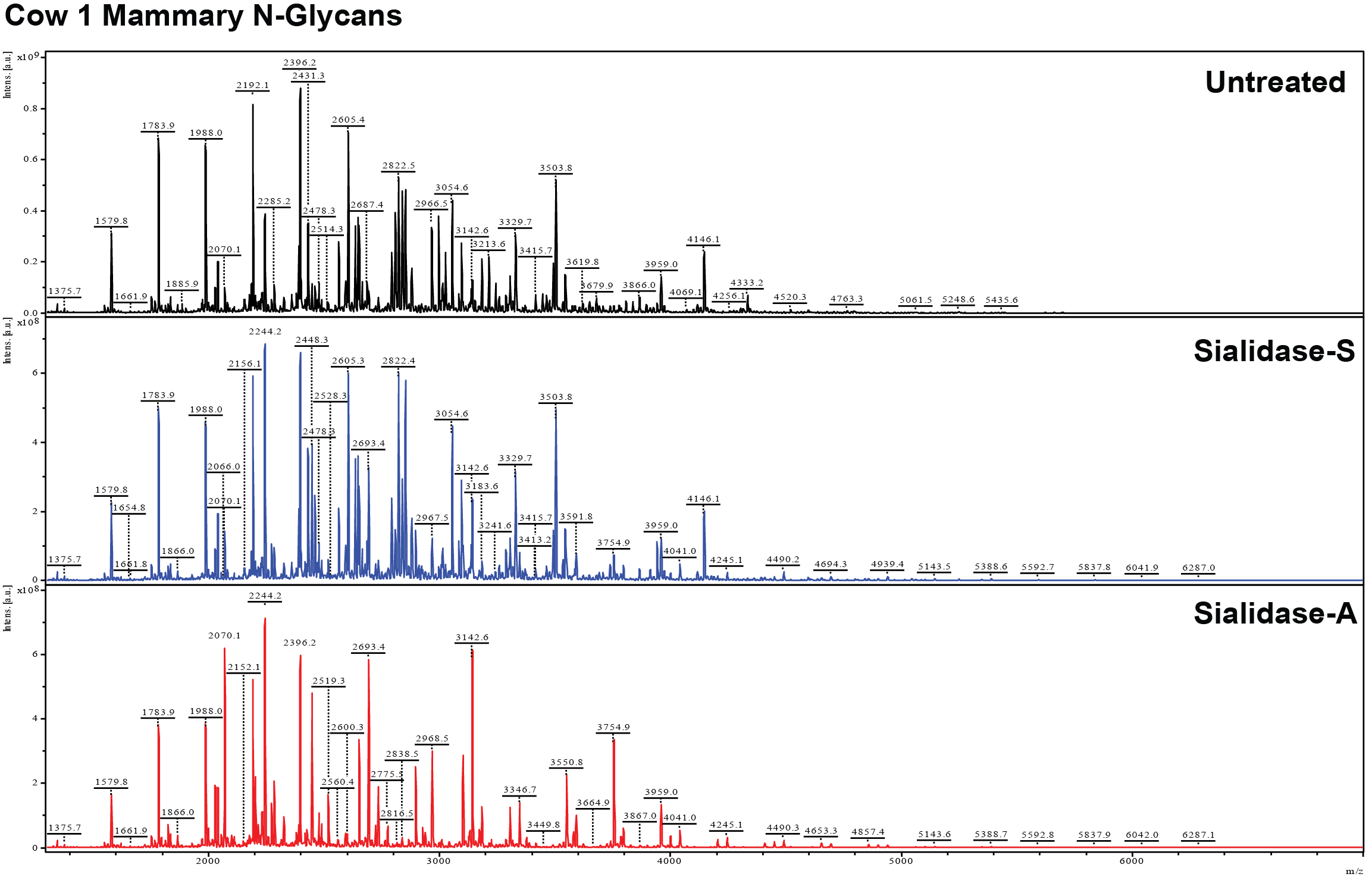

**Supplementary Figure S6: Annotated MALDI-TOF spectra of bovine mammary N-glycans from cow 1 (lactating).**

Bovine mammary N-glycans A) untreated (black), B) treated with sialidase-S (blue), or C) treated with sialidase-A (red). Peak annotation conducted by flexAnalysis 5.0 software.

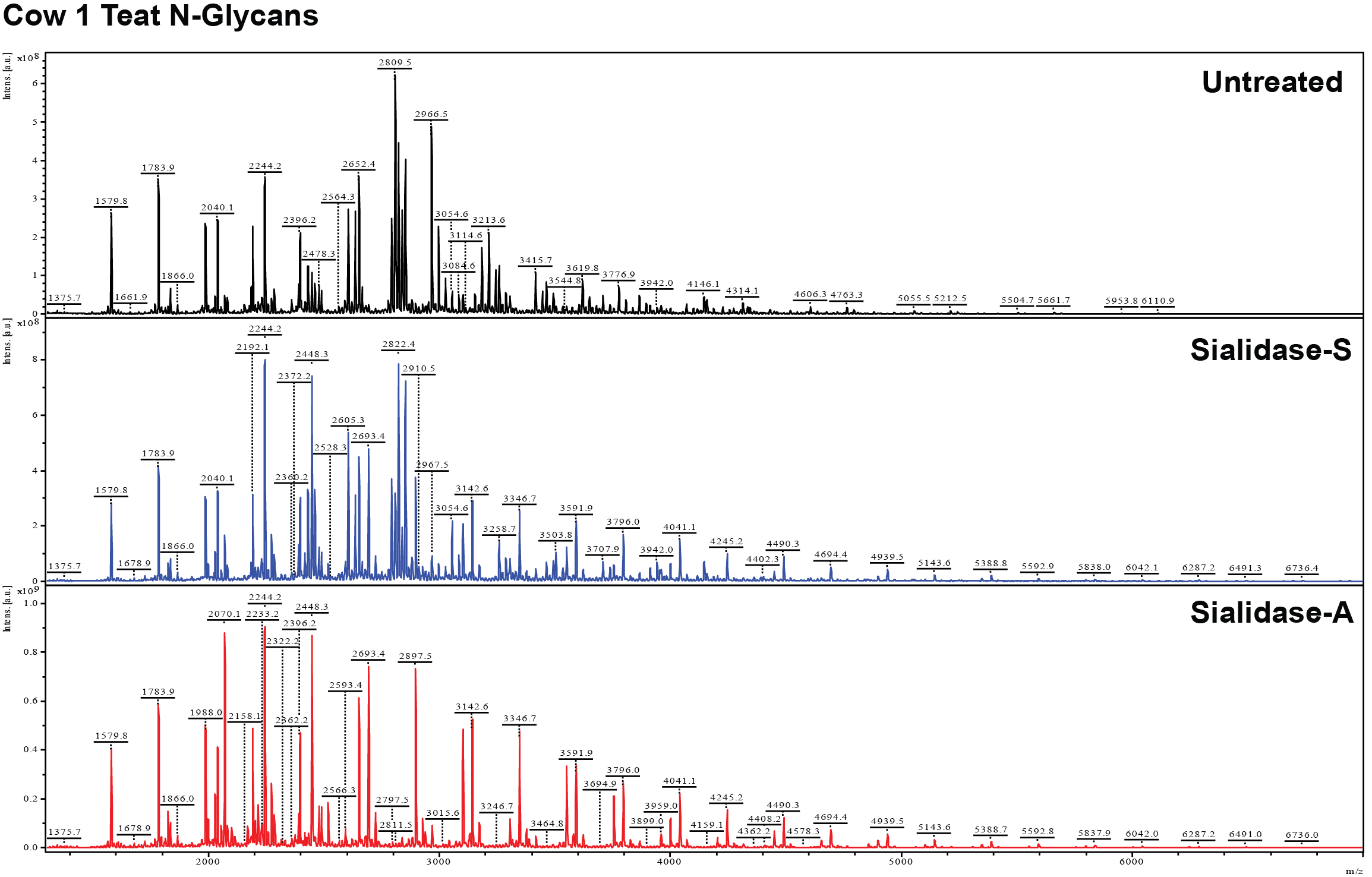

**Supplementary Figure S7: Annotated MALDI-TOF spectra of bovine teat N-glycans from cow 1 (lactating).**

Bovine teat N-glycans A) untreated (black), B) treated with sialidase-S (blue), or C) treated with sialidase-A (red). Peak annotation conducted by flexAnalysis 5.0 software.

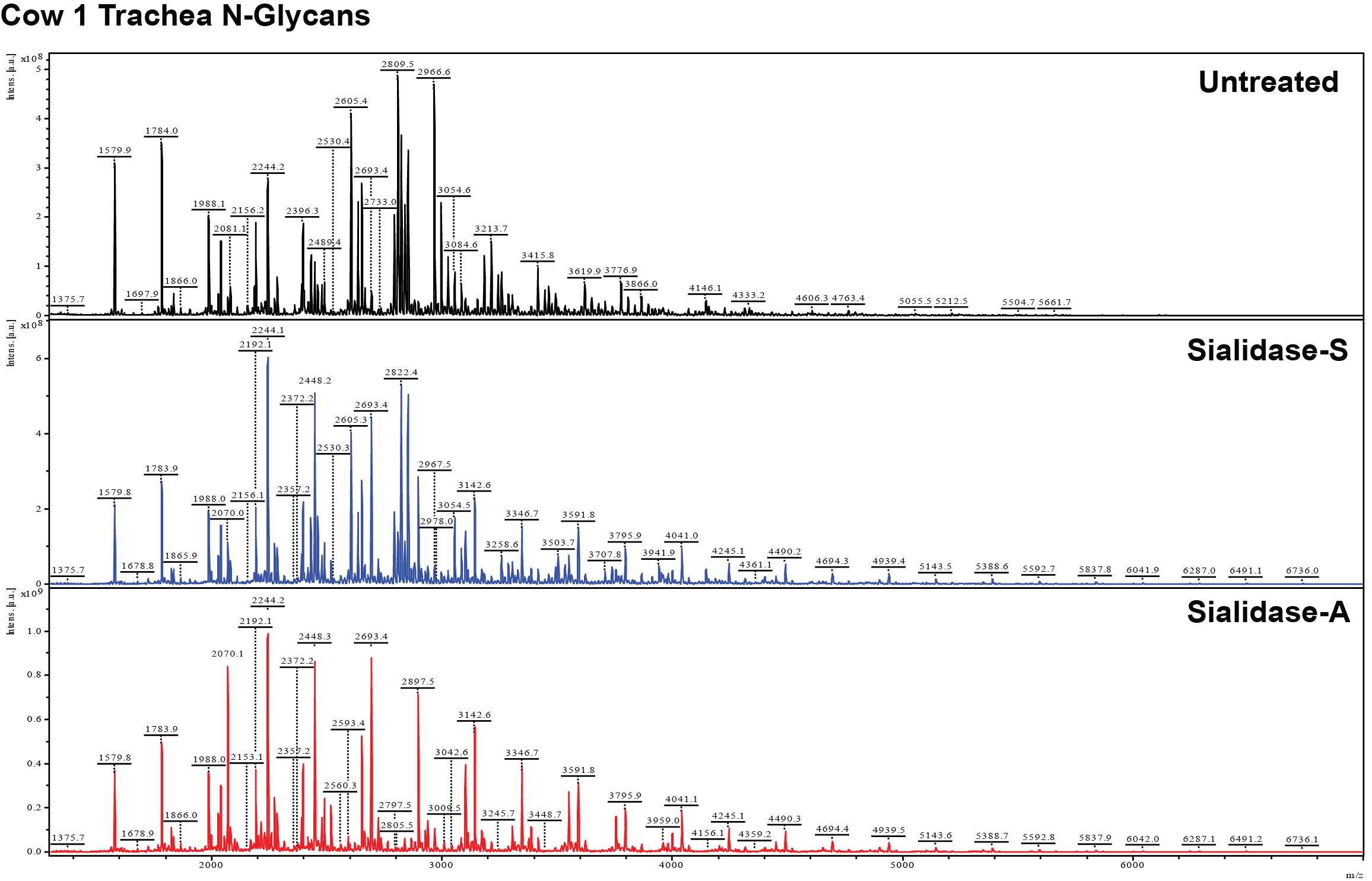

**Supplementary Figure S8: Annotated MALDI-TOF spectra of bovine trachea N-glycans from cow 1 (lactating).**

Bovine trachea N-glycans A) untreated (black), B) treated with sialidase-S (blue), or C) treated with sialidase-A (red). Peak annotation conducted by flexAnalysis 5.0 software.

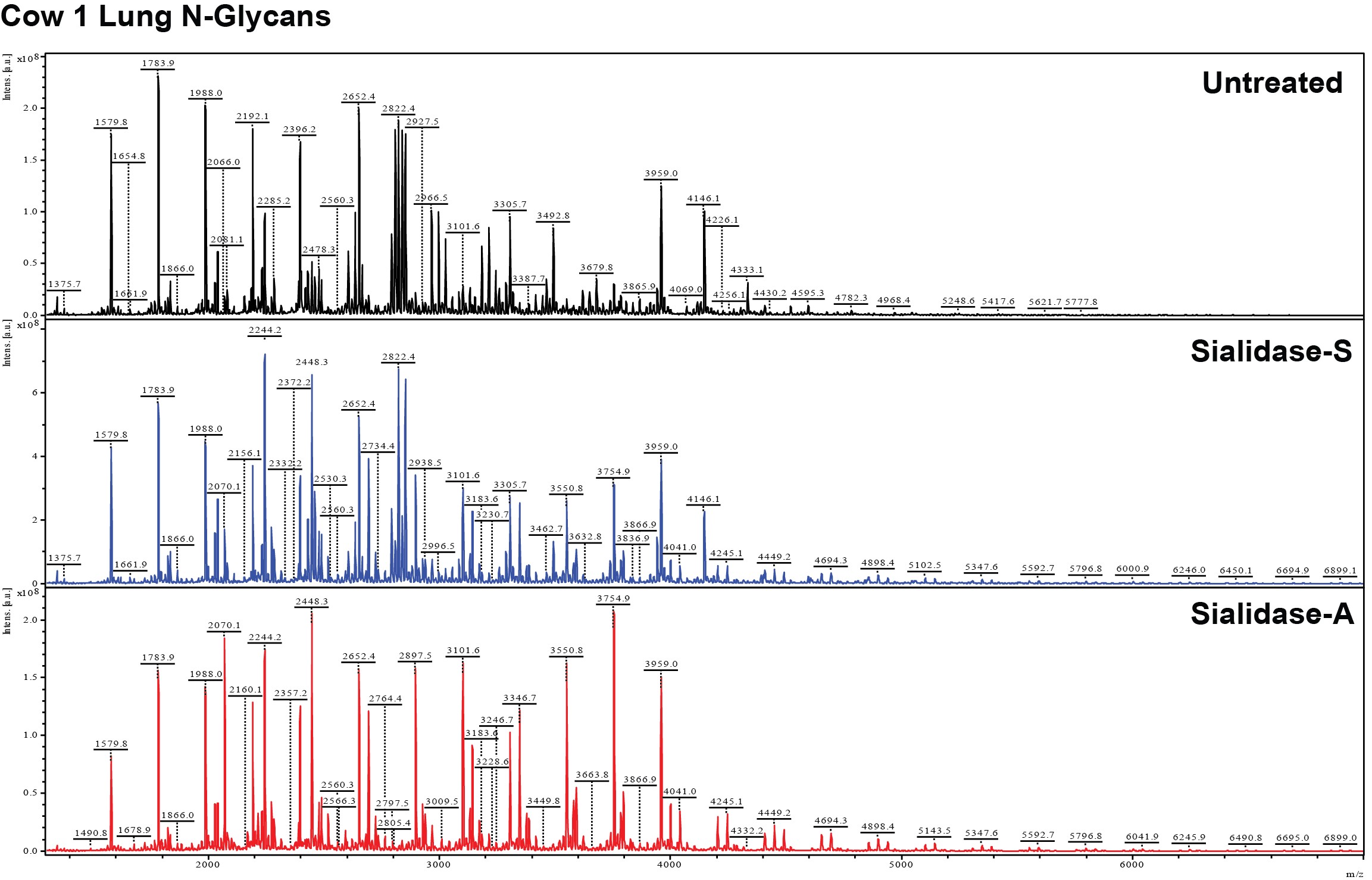

**Supplementary Figure S9: Annotated MALDI-TOF spectra of bovine lung N-glycans from cow 1 (lactating).**

Bovine lung N-glycans A) untreated (black), B) treated with sialidase-S (blue), or C) treated with sialidase-A (red). Peak annotation conducted by flexAnalysis 5.0 software.

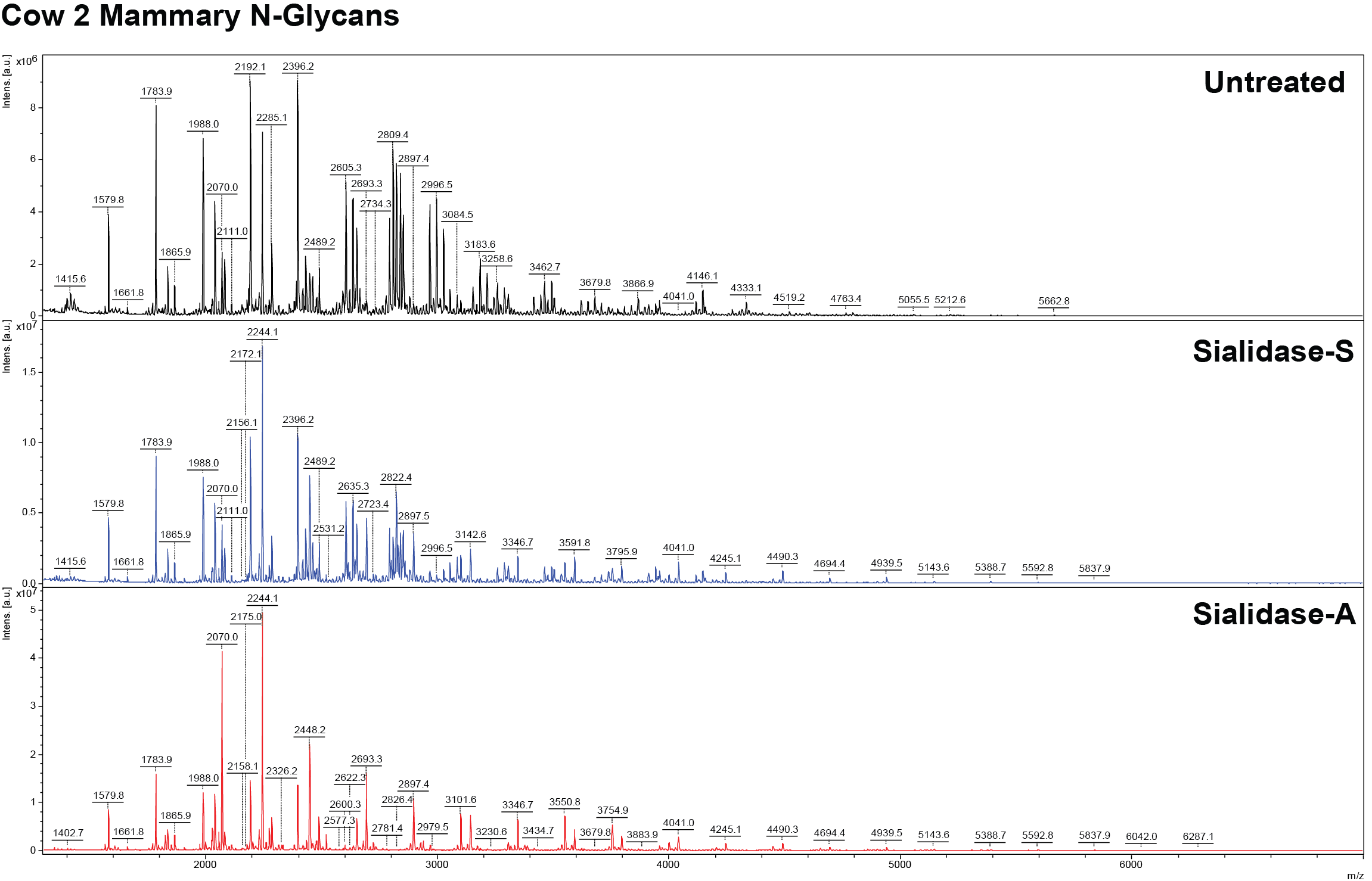

**Supplementary Figure S10: Annotated MALDI-TOF spectra of bovine mammary N-glycans from cow 2 (non-lactating).**

Bovine mammary N-glycans A) untreated (black), B) treated with sialidase-S (blue), or C) treated with sialidase-A (red). Peak annotation conducted by flexAnalysis 5.0 software.

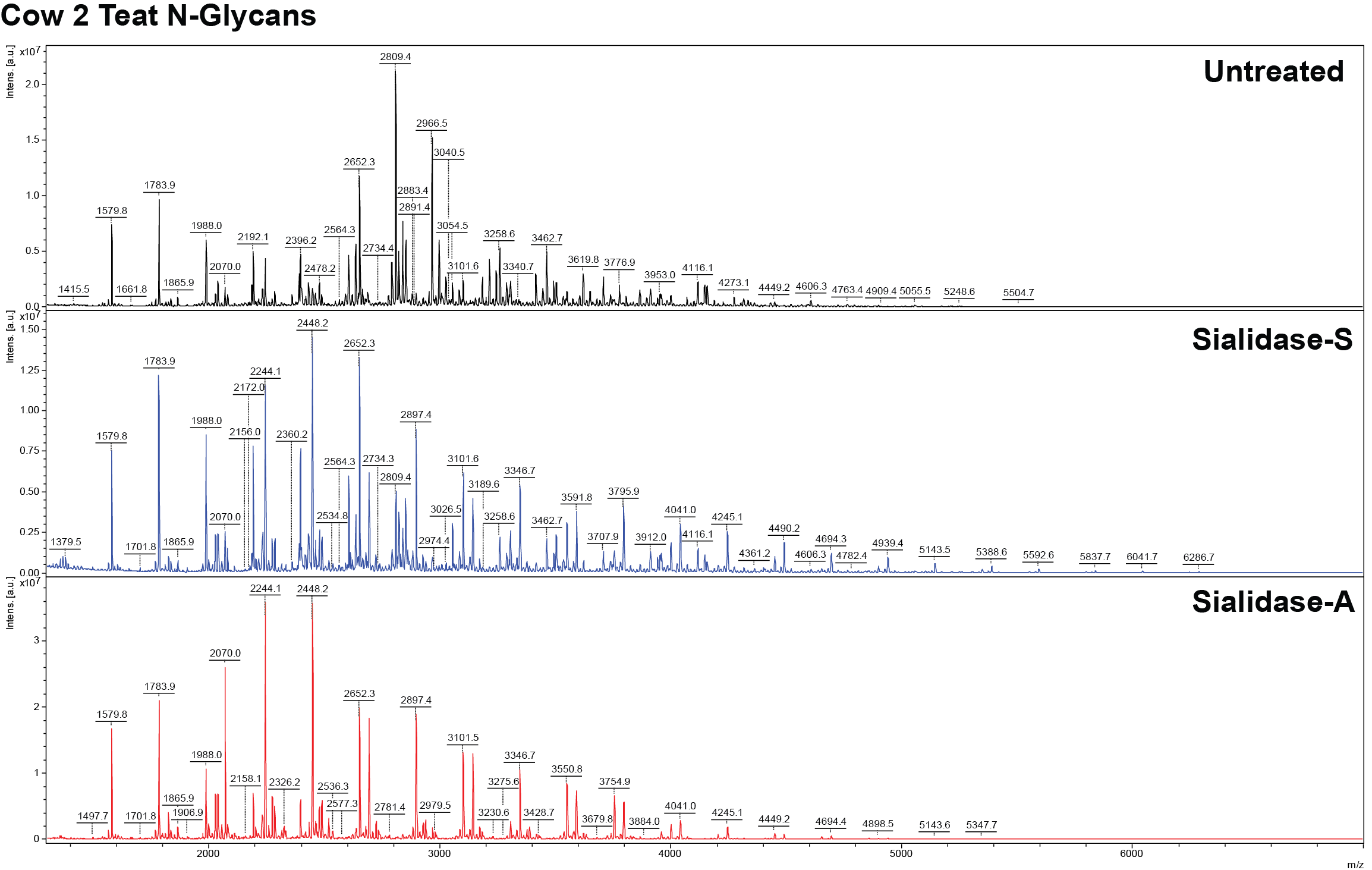

**Supplementary Figure S11: Annotated MALDI-TOF spectra of bovine teat N-glycans from cow 2 (non-lactating).**

Bovine teat N-glycans A) untreated (black), B) treated with sialidase-S (blue), or C) treated with sialidase-A (red). Peak annotation conducted by flexAnalysis 5.0 software.

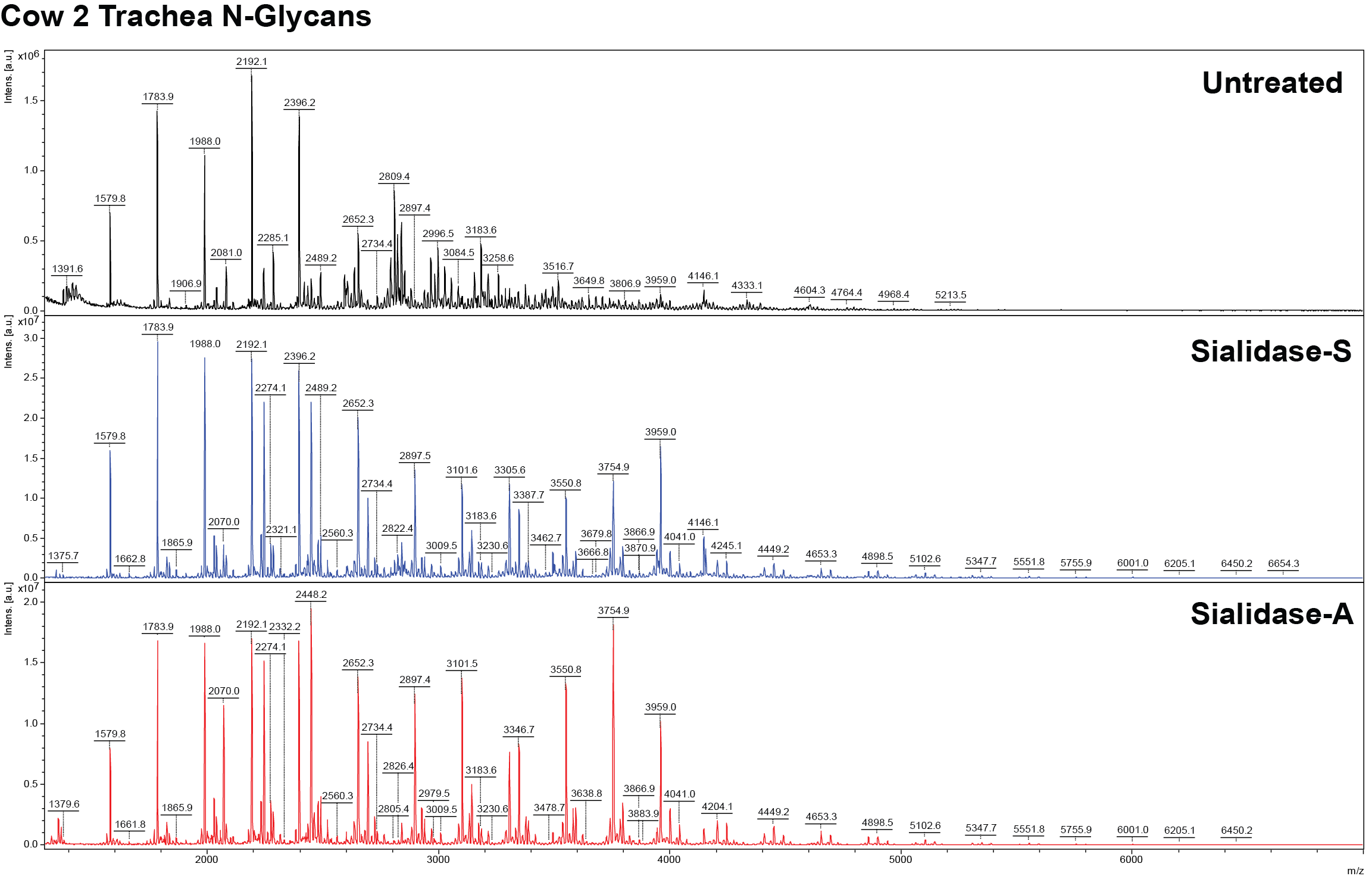

**Supplementary Figure S12: Annotated MALDI-TOF spectra of bovine trachea N-glycans from cow 2 (non-lactating).**

Bovine trachea N-glycans A) untreated (black), B) treated with sialidase-S (blue), or C) treated with sialidase-A (red). Peak annotation conducted by flexAnalysis 5.0 software.

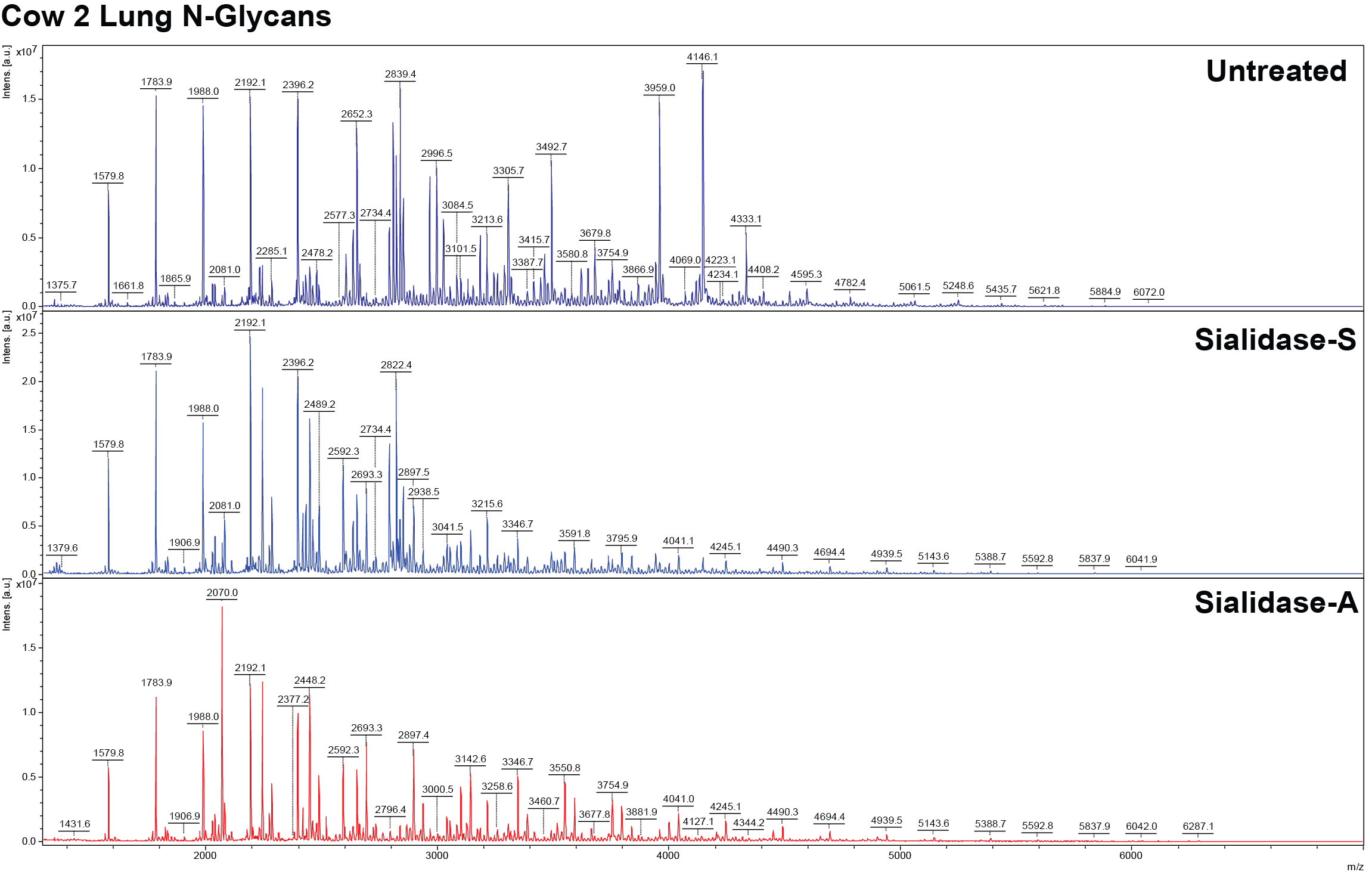

**Supplementary Figure S13: Annotated MALDI-TOF spectra of bovine lung N-glycans from cow 2 (non-lactating).**

Bovine lung N-glycans A) untreated (black), B) treated with sialidase-S (blue), or C) treated with sialidase-A (red). Peak annotation conducted by flexAnalysis 5.0 software.

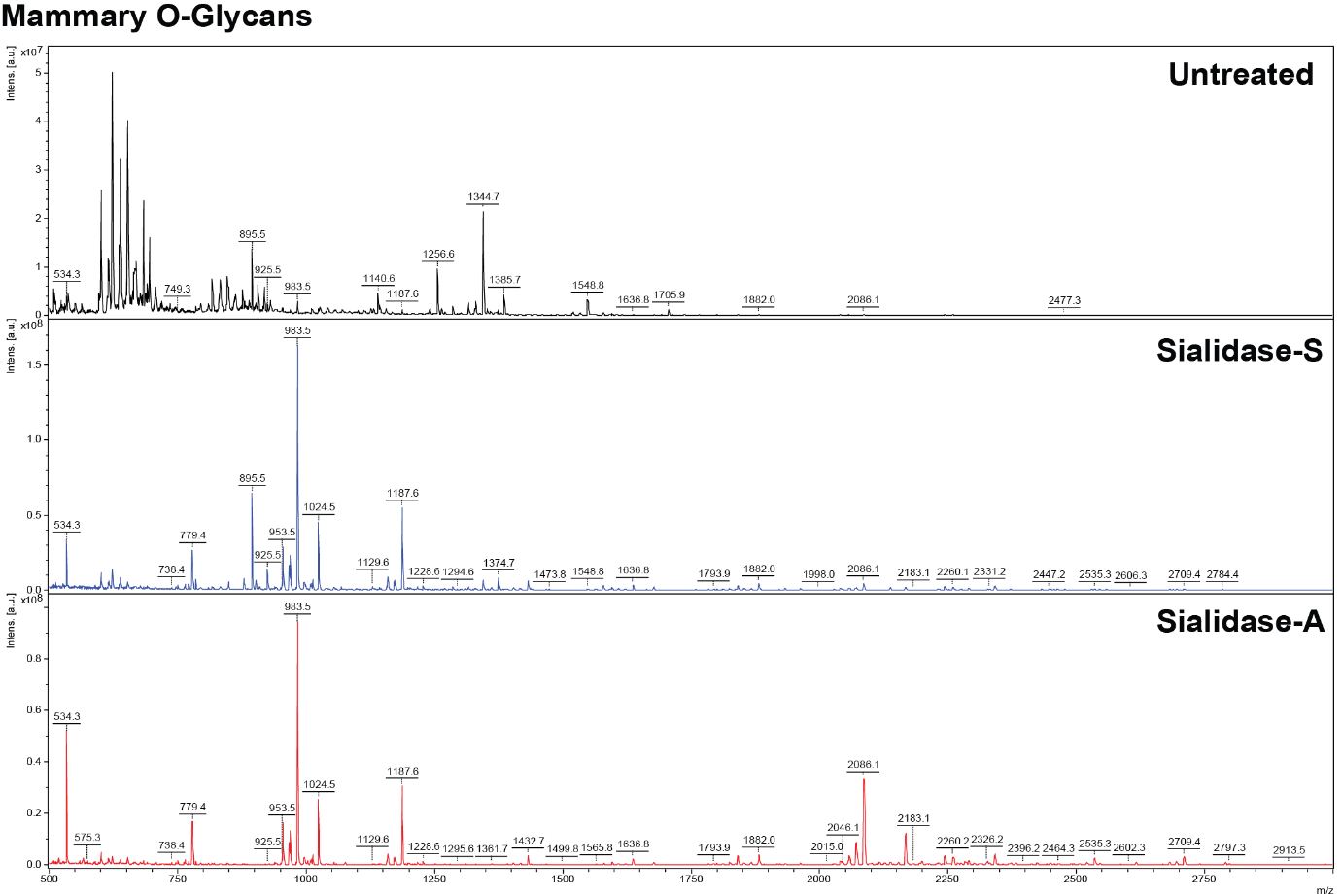

**Supplementary Figure S14: Annotated MALDI-TOF spectra of bovine mammary O-glycans from cow 2 (non-lactating).**

Bovine mammary O-glycans A) untreated (black), B) treated with sialidase-S (blue), or C) treated with sialidase-A (red). Peak annotation conducted by flexAnalysis 5.0 software.

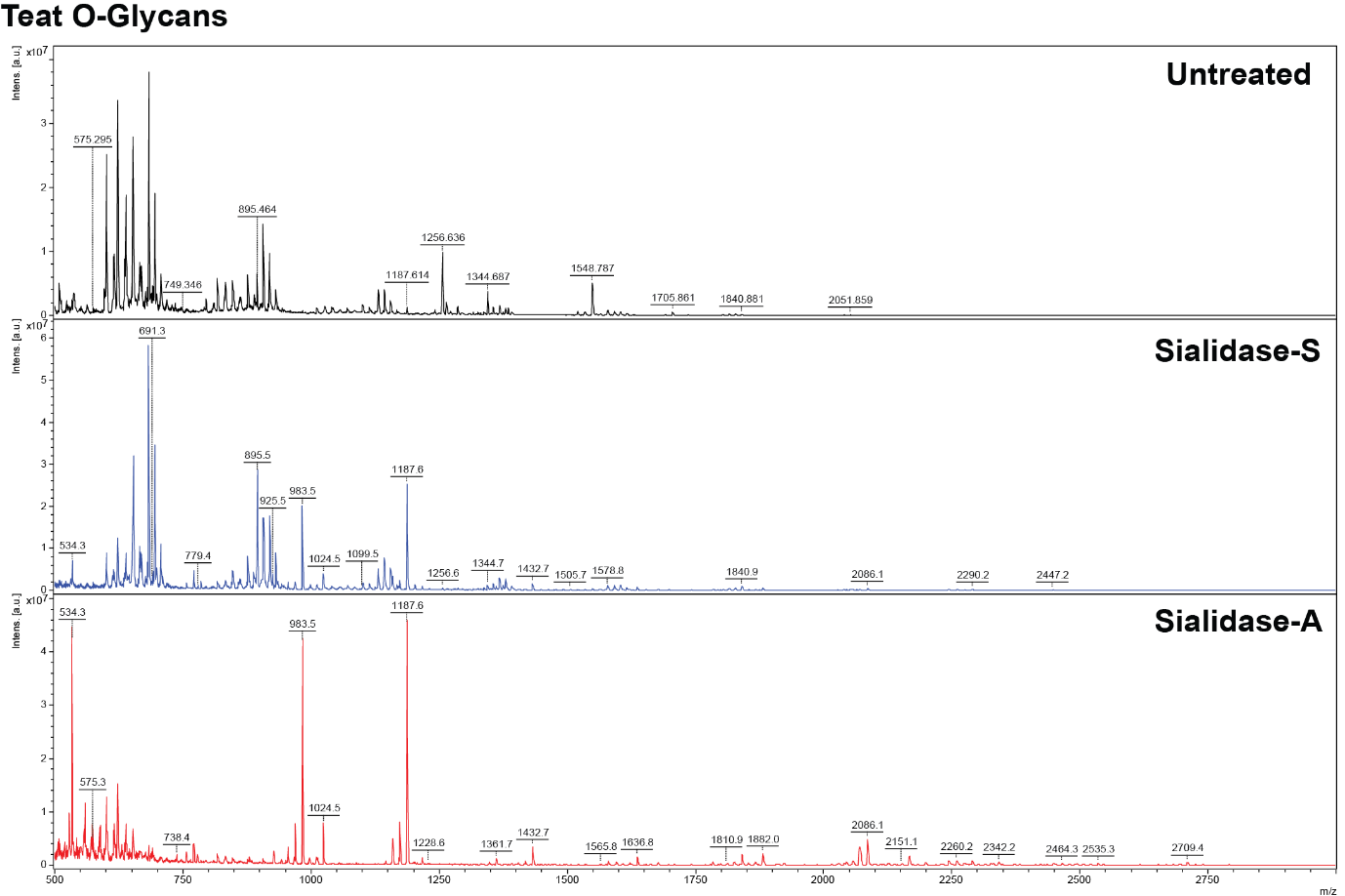

**Supplementary Figure S15: Annotated MALDI-TOF spectra of bovine teat O-glycans from cow 2 (non-lactating).**

Bovine teat O-glycans A) untreated (black), B) treated with sialidase-S (blue), or C) treated with sialidase-A (red). Peak annotation conducted by flexAnalysis 5.0 software.

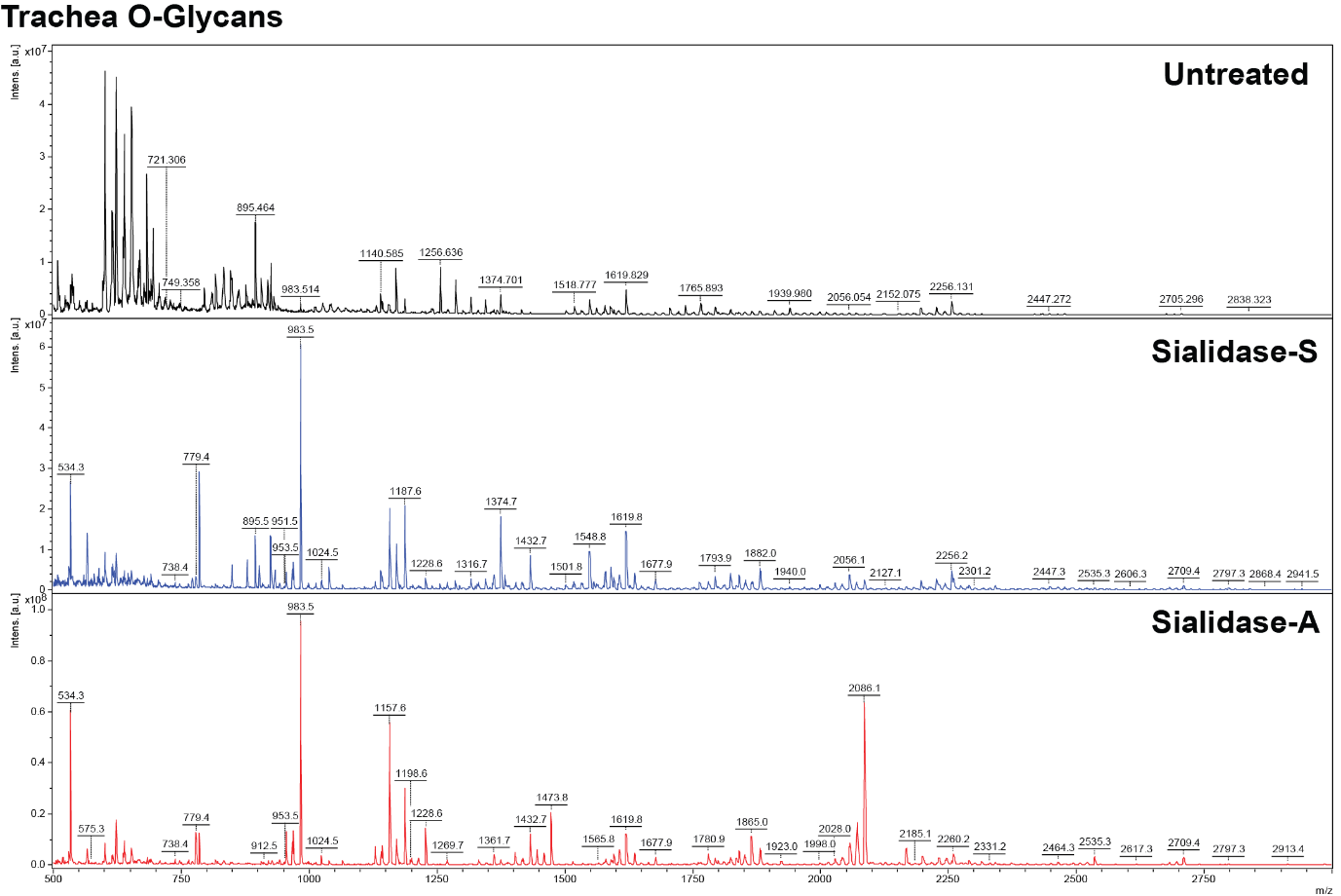

**Supplementary Figure S16: Annotated MALDI-TOF spectra of bovine trachea O-glycans from cow 2 (non-lactating).**

Bovine trachea O-glycans A) untreated (black), B) treated with sialidase-S (blue), or C) treated with sialidase-A (red). Peak annotation conducted by flexAnalysis 5.0 software.

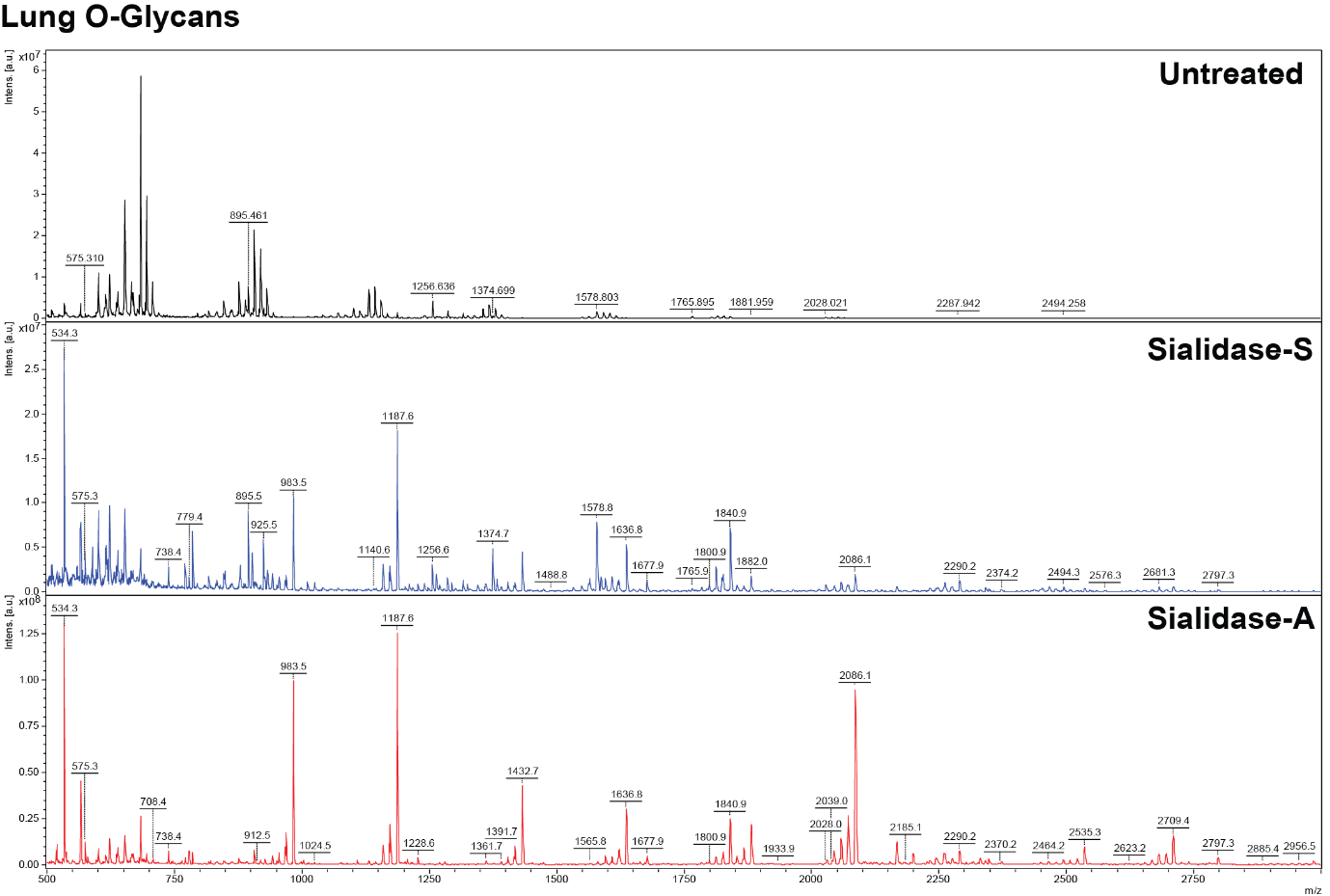

**Supplementary Figure S17: Annotated MALDI-TOF spectra of bovine lung O-glycans from cow 2 (non-lactating).**

Bovine lung O-glycans A) untreated (black), B) treated with sialidase-S (blue), or C) treated with sialidase-A (red). Peak annotation conducted by flexAnalysis 5.0 software.

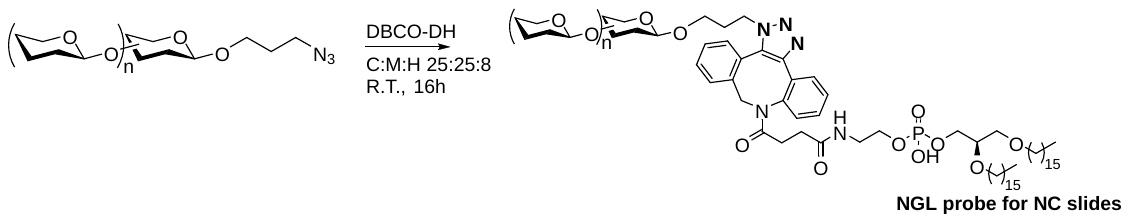

**Supplementary Figure S18. Conversion of azido glycans into NGLs.**

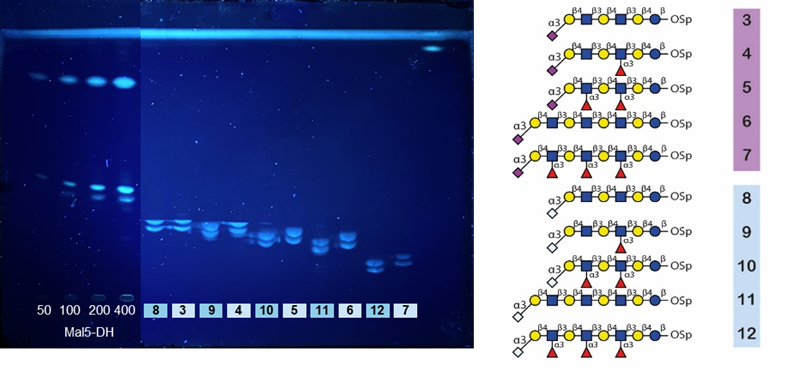

**Supplementary Figure S19. HPTLC quantitation of generated NGLs.** Maltopentaose NGL standards (50, 100, 200, and 400 pmol; lanes 1–4) were applied and the HPTLC plate was developed in a solvent system of CHCl₃:MeOH:H₂O (60:35:8, v/v/v). Quantification of the generated NGLs was performed based on UV absorbance measurements of the primulin-stained bands on the developed plates. Double bands observed correspond to the two isomeric forms of the azido–DBCO linkage. Quantification was performed based on the combined intensity of these two isomeric bands.

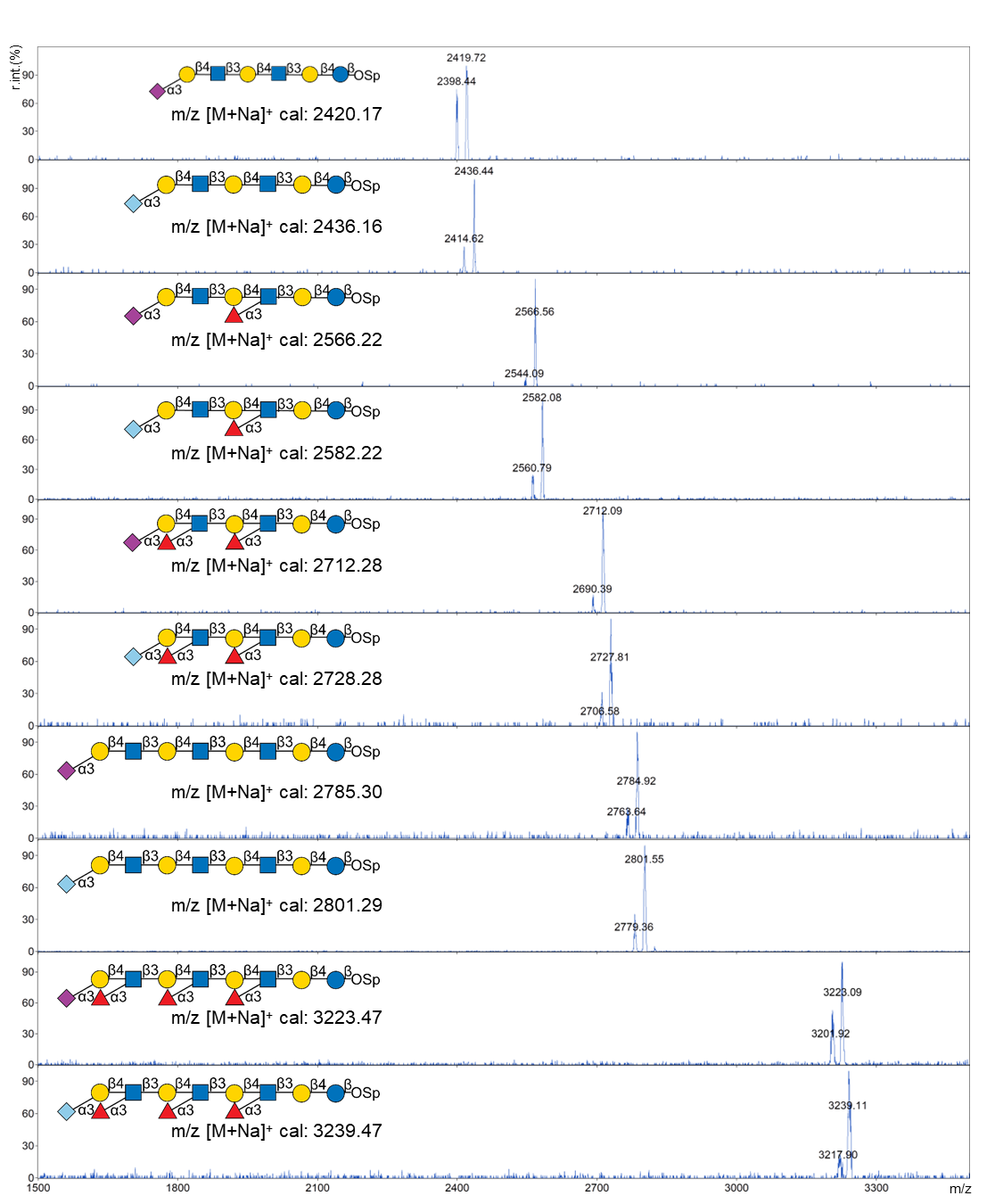
**Supplementary Figure S20. MS spectra of synthesized NGLs.** Mass spectrometric analysis was performed in positive ion mode, revealing characteristic signals corresponding to [M+H]⁺ and [M+Na]⁺ ions.

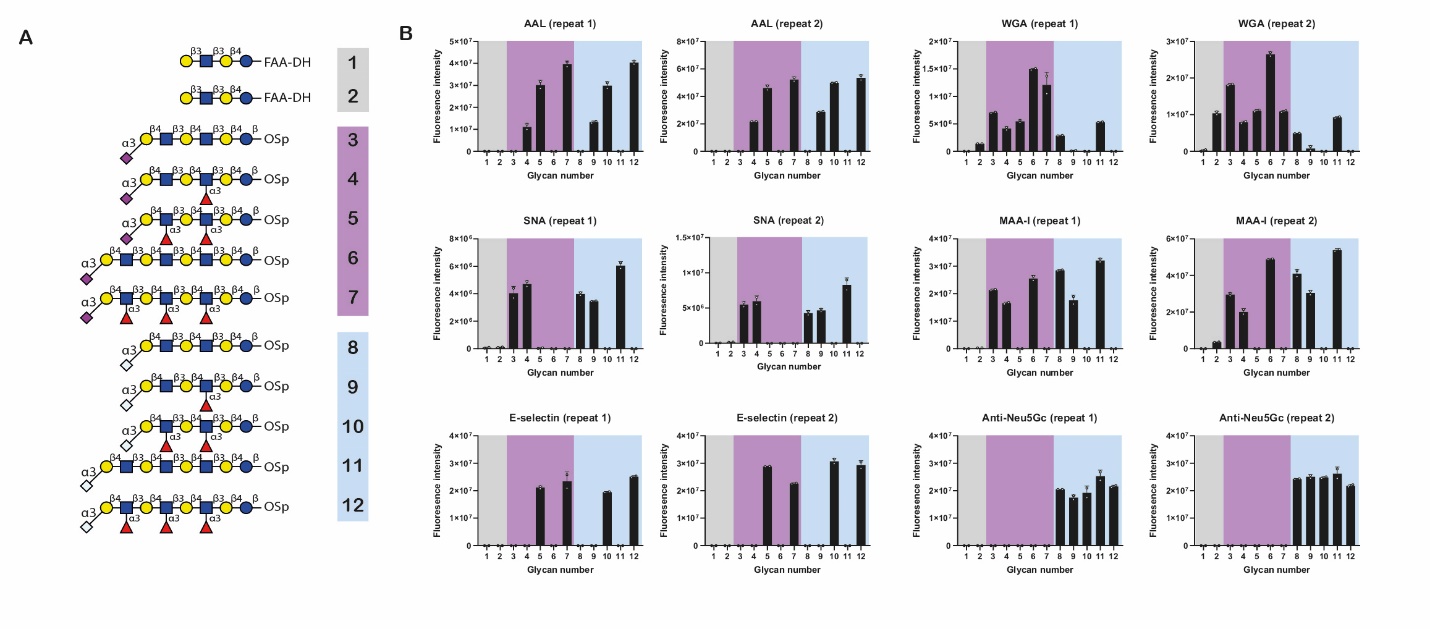

**Supplementary Figure S21. Quality control of the glycan array using lectins and antibody.** Binding of lectins and antibodies to a panel of receptor analogues immobilized to glycan arrays. Non-sialylated analogues backed in grey, analogues containing NeuAc backed in pink and NeuGc backed in pale blue. N = 2 biological repeats shown with N = 2 technical repeats.

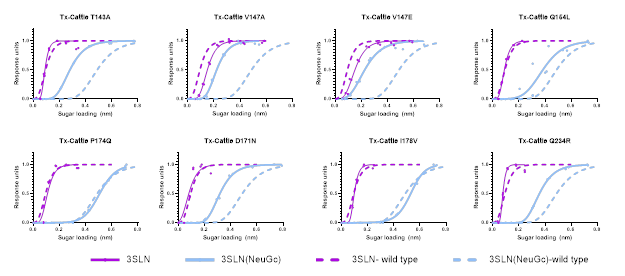

**Supplementary Figure S22. Additional bio-layer interferometry data of cattle H5N1 mutants.** Bio-layer interferometry plots showing binding of purified viruses to different receptor analogues shown in panel**.** Solid (mutant) and dashed (wildtype), pink (3SLN-NeuAc) and pale blue (3SLN-NeuGc) lines indicate receptor binding to analogues. Data points shown as duplicate repeats with a asymmetric Sigmoidal 5PL curve fitted, constrained to 0 and 1 at the top and bottom.

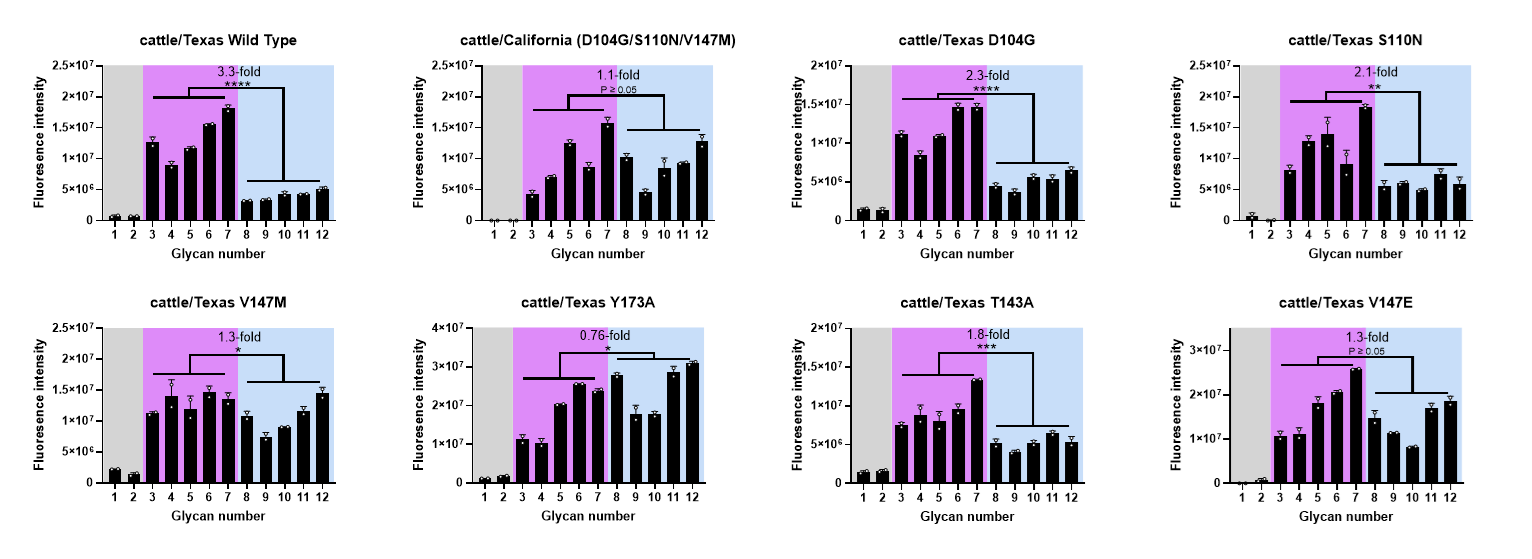

**Supplementary Figure S23. Extended glycan array data of cattle H5N1 mutants.** Binding of purified viruses to a panel of receptor analogues immobilized to glycan arrays. Non-sialylated analogues backed in grey, analogues containing NeuAc backed in pink and NeuGc backed in pale blue. Data shown as N = 2 technical duplicates (second biological replicate shown in supplementary figure S18). Statistics performed by paired, two-tailed t-tests with individual variance for each group – e.g. comparing each matched set of analogues containing either NeuAc or NeuGc.

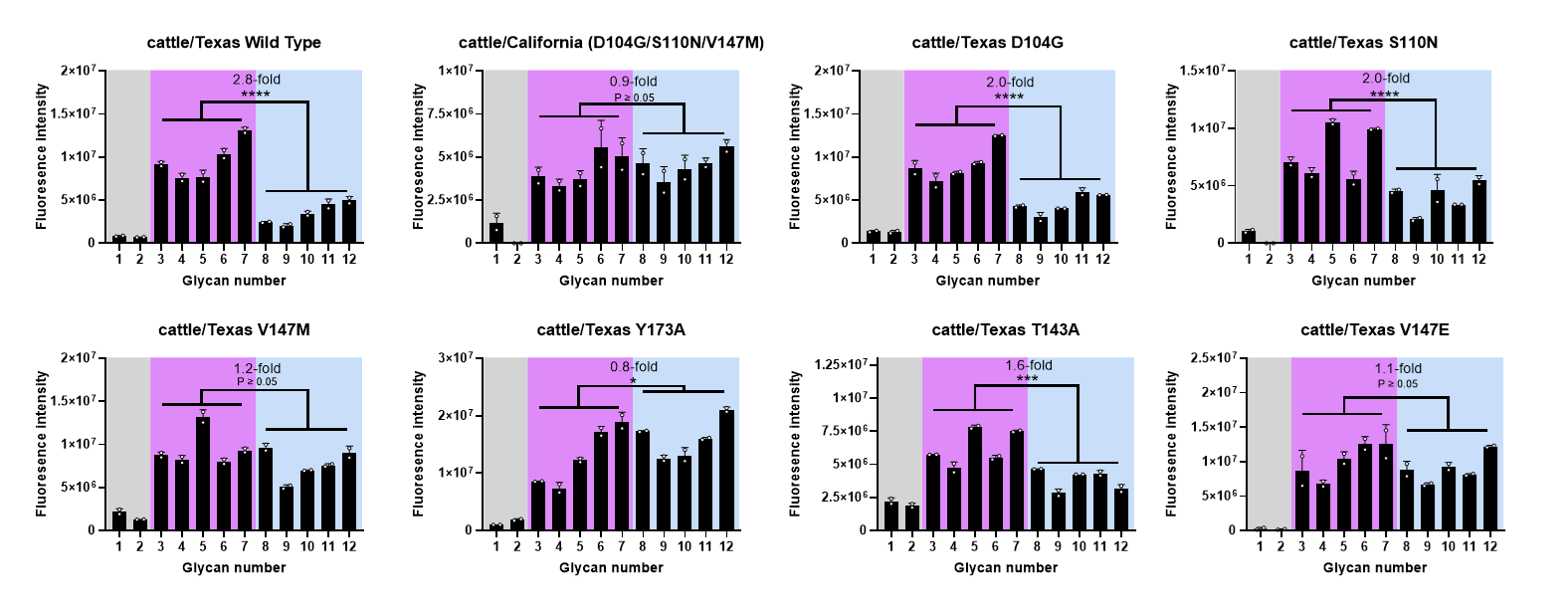

**Supplementary Figure S24. Biological repeat of glycan array data.** Binding of purified viruses to a panel of receptor analogues immobilized to glycan arrays. Non-sialylated analogues backed in grey, analogues containing NeuAc backed in pink and NeuGc backed in pale blue. Data shown as N = 2 technical duplicates (first biological replicate shown in supplementary figure S17). Statistics performed by paired, two-tailed t-tests with individual variance for each group – e.g. comparing each matched set of analogues containing either NeuAc or NeuGc.

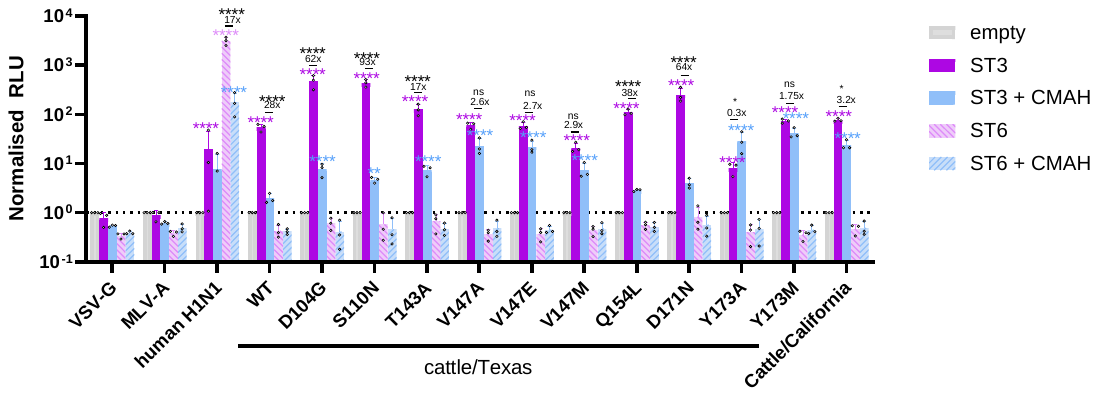

**Supplementary Figure S25. Extended pseudovirus entry data into 293 cells knocked out for sialyl-transferase expression.** Relative entry of pseudoviruses into 293 cells that have been genetically ablated for sialyl-transferases, and then transiently transfected with either empty vector, an expression plasmid for sialyl-transferases (ST3GAL4 or ST6GAL1) with or without chimpanzee CMAH. Data plotted as mean + S.D. with the mean values from N = 3 independent repeats plotted. Statistics performed using 2-way ANOVA with multiple comparisons upon log-transformed data. Only differences above empty vector, and between ST3GAL4 and ST3GAL4+CMAH shown. Log-normality throughout determined by Shapiro-Wilk test and QQ plot. Significance shown by asterisks indicating: *, 0.05 ≥ P > 0.01; **, 0.01 ≥ P > 0.001; ***, 0.001 ≥ P > 0.0001; ****, P ≤ 0.0001.

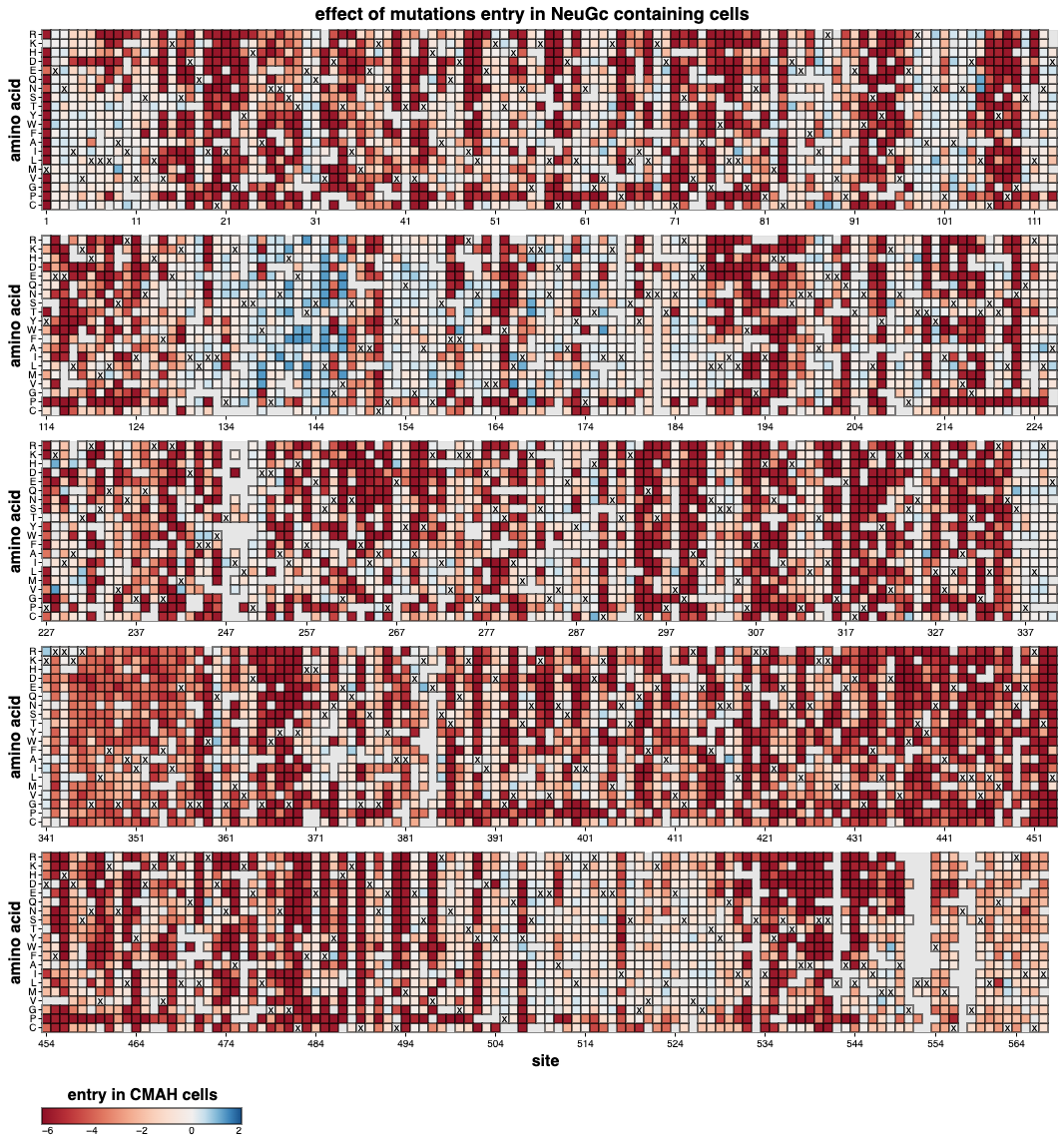

**Supplementary Figure S26. Whole HA deep mutational scanning heatmap of entry into 293-CMAH cells.** Interactive heatmap available at <https://dms-vep.org/Flu_H5_American-Wigeon_South-Carolina_2021-H5N1_DMS_NeuGc/htmls/entry_in_293-CMAH_entry_wrapped_heatmap.html> .

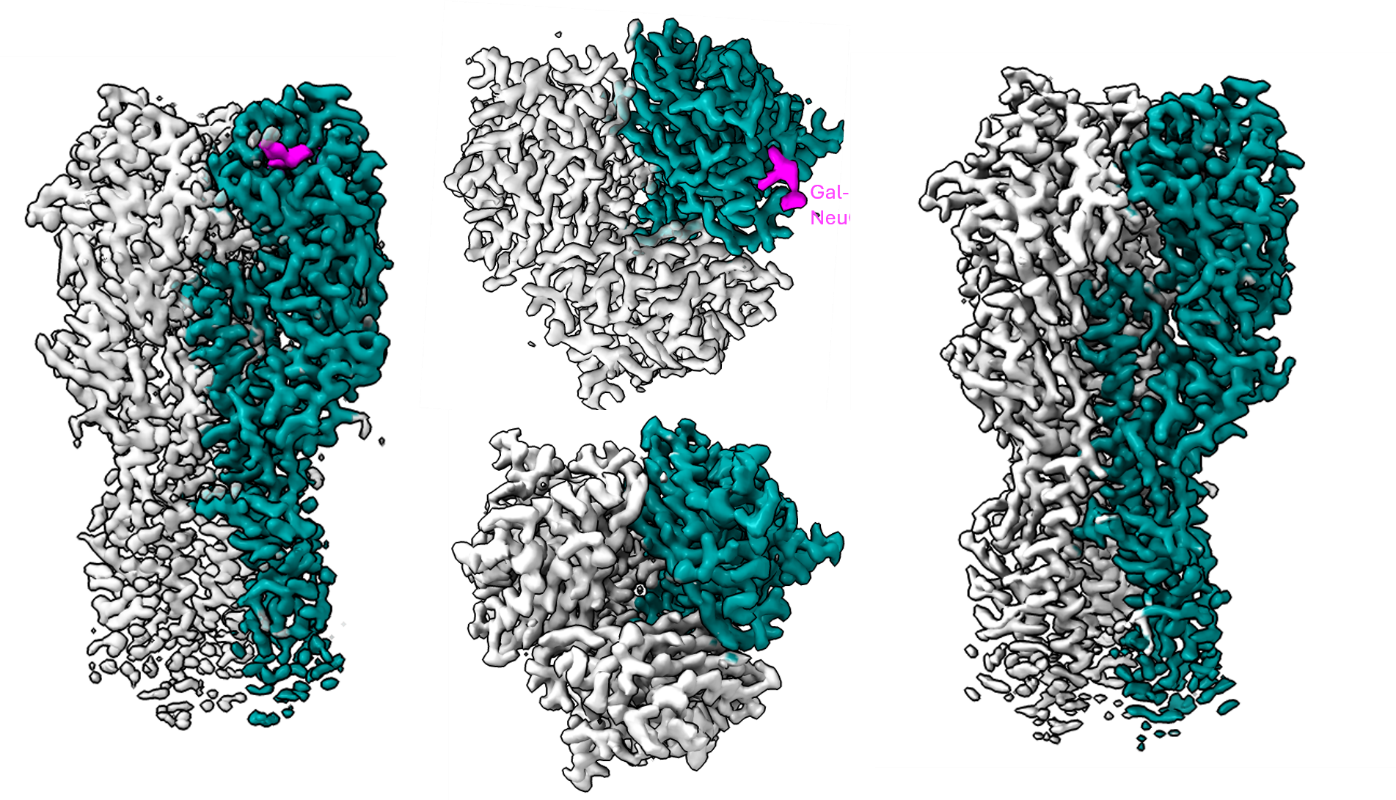

**Supplementary Figure S27. Density maps of the HA of cattle/California bound to the NeuGc containing receptors.**

**Supplementary Tables**

**Supplementary Table 2. HA numbering conversion table.**

| **Immature H5 numbering** | **Mutation** | **Mature H5 numbering** | **Equivalent H3 numbering** |
| --- | --- | --- | --- |
| 104 | D104G | 88 | 95 |
| 110 | S110N | 94 | 101 |
| 130 | I130T | 114 | 121 |
| 133 | I133F | 117 | 124 |
| 136 | S136N | 120 | 125b |
| 140 |  | 124 | 129 |
| 141 | H141K | 125 | 130 |
| 142 | E142W | 126 | 131 |
| 143 | T143A/F | 127 | 132 |
| 145 | L145Q | 129 | 133a |
| 147 | V147M/E/A | 131 | 135 |
| 154 | Q154L | 138 | 142 |
| 168 | K168G | 152 | 156 |
| 169 | K169R | 153 | 157 |
| 171 | D171G/N | 155 | 159 |
| 172 | A172T | 156 | 160 |
| 173 | Y173A/M | 157 | 161 |
| 336 | S336N | 320 | 323 |

**Supplementary Table 6. GBPs used in glycan array.**

| GBPs | Source | Catalogue # | Used conc |
| --- | --- | --- | --- |
| Aleuria Aurantia Lectin (AAL) | Vector | B-1395 | 20 μg/mL |
| Wheat Germ Agglutinin (WGA) | Vector | B-1025 | 20 μg/mL |
| Sambucus Nigra Lectin (SNA) | Vector | B-1305 | 50 μg/mL |
| Maackia Amurensis Lectin I (MAA-I) | Vector | B-1315 | 100 μg/mL |
| Anti-Neu5Gc Poly21469 (Chicken IgY) | Biolegend | 146903 | 1:200 dilution |
| Anti-CK IgY Polyclonal (biotinylated) | Abcam | ab207998 | 1:200 dilution |
| Murine E-Selectin (Human IgM)* | John B. Lowe, University of Michigan Medical School^57^ | | 1:4 dilution |
| Anti-Human IgM (biotinylated) | Vector | BA-3020 | 1:200 dilution |
| Anti TX-Cattle WT chicken polyserum |  |  | 1:100 dilution |

** E-Selectin was kindly provided by John B. Lowe, University of Michigan Medical School.*

Supplementary Table 7. MIRAGE compliant Glycan Microarray Document

Supplementary Glycan Microarray Document Based on MIRAGE Guidelines (doi:10.3762/mirage.3)

|  | Description |
| --- | --- |
| 1. Sample: Glycan Binding Sample | |
| Description of Sample | The anti-carbohydrate antibodies, biotinylated plant lectins and immune lectin receptors used in this study are detailed in **Supplementary Table 6**. |
| Sample modifications | Not relevant. |
| Assay protocol | Microarray binding analyses were performed at R.T. essentially as described^58^. In brief, the NC slides were treated with blocking buffer for 1 h prior to incubations. The overlay steps of the Tx virus proteins were performed in blocking buffer A containing 0.2%BSA, 100 μM of Oseltamivir and Zanamivir in HEPES-Buffered Saline (HBS) (5 mM HEPES buffer pH 7.4, 150 mM NaCl), whereas the incubations of plant lectins and anti-carbohydrate antibodies were performed in blocking buffer B containing 1%BSA, 10 mM CaCl_2_ in HBS. The concentration of the antibodies and lectins are indicated in **Supplementary Table 6**. Stepwise incubations were performed for 1 h per overlay step, except for virus incubation for 1.5 h followed with fixing using 4% formaldehyde at 0^o^C for 0.5 h. The final detection step was carried out using Alexa Fluor-647-labeled streptavidin (1 μg/mL, Molecular Probes, Eugene) for 0.5 h. |
| 2. Glycan Library | |
| Glycan description for defined glycans | The sequence information on the NGL probes are in **Supplementary Figure 15.** |
| Glycan description for undefined glycans | Not relevant. |
| Glycan modifications | The synthesis of NGL probes were described above under Supplementary Methods. |
| 3. Printing Surface; e.g., Microarray Slide | |
| Description of surface | For noncovalent array, UniSart® Microarray slide with 16 nitrocellulose membrane pads from Sartorius (Stedim, Goettingen, Germany) were used. |
| Manufacturer | Indicated above. |
| Custom preparation of surface | Not relevant. |
| Noncovalent Immobilization | The lipid-linked oligosaccharide probes were formulated as liposomes by adding carrier lipids, 1,2-dihexanoyl-sn-glycero-3-phosphocholine (DHPC) and cholesterol (both from Merck) for arraying and non-covalent immobilization on nitrocellulose-coated glass slides essentially as described^58^. |
| Covalent Immobilization | Not relevant. |
| 4. Arrayer (Printer) | |
| Description of Arrayer | The printing of the microarrays was performed with Nano-Plotter from Gesim, Germany. |
| Dispensing mechanism | Non-contact liquid delivery with four dispensing tips. |
| Glycan deposition | For noncovalent printing, single droplet (approximate 0.33 nL) was dispensed for each spot. |
| Printing conditions | For noncovalent arrays, the ‘liposome’ printing solutions (aqueous based) contained 100 pmol/μL of DHPC and cholesterol as lipid carriers in addition to the lipid-linked glycan probes. The concentrations of the lipid-linked glycan probes were 5 and 15 pmol/μL for the 2 and 5 fmol per spot levels, respectively. The printing solutions also contained Cy3 NHS ester (GE Healthcare) at 20 ng/ml as a marker to monitor the printing process. The resulting arrays were printed at R.T. under 58% relative humidity. |
| 5. Glycan Microarray with “Map” | |
| Array layout | For the noncovalent arrays, each slide contained 16 pads in an 8 × 2 format. Each subarray included 12 NGLs, each printed at two concentrations (2 and 5 fmol per spot) in duplicate (resulting in four spots per glycan in a row). |
| Glycan identification and quality control | The details of glycan sequences are included in **Supplementary Figure 15**. Quality control with sequence-specific lectins and anti-carbohydrate antibodies were listed in **Supplementary Fig. 15**. |
| 6. Detector and Data Processing | |
| Scanning hardware | GenePix 4300A (Molecular Devices, Berkshire, UK) |
| Scanner settings | The scanning of the microarray slides after incubation and drying was carried out at a resolution of 5 µm/pixel, using the red laser channel (scan wavelength 635 nm).  For noncovalent NC slides PMT Voltages were set at 350V and the scan power ranging from 15% to 100% was used to avoid signal saturation. |
| Image analysis software | GenePix® Pro 7 (Molecular Devices, Berkshire, UK) |
| Data processing | The gpr files were processed in pre-set Excel templates. No particular normalization method or statistical analysis was used. |
| 7. Glycan Microarray Data Presentation | |
| Data presentation | Binding results are presented as histogram charts in the main text **Fig. 3** and additional figures in **Supplementary Fig. 15,** **17 and 18.**  Binding score data are shown in **Supplementary Tables 6** and **7**. |
| 8. Interpretation and Conclusion from Microarray Data | |
| Data interpretation | No software or algorithms were used to interpret processed data. |
| Conclusions | The conclusion of the findings of the Tx virus binding are in the discussion section. |
